# IL-1β engages distinct peripheral sensory circuits to suppress feeding across time

**DOI:** 10.64898/2026.02.23.707454

**Authors:** Nikolas W. Hayes, Jessica L. Xia, Joshua A. Frydman, Marcos J. Duran, Kamil K. Gebis, Carolyn M. Lorch, Haley S. Province, Esther Tang, Lisa R. Beutler

## Abstract

Loss of appetite is a hallmark of acute and chronic inflammation, and sustained anorexia causes malnutrition and worsens disease outcomes. Interleukin-1β (IL-1β) is one of the most potently anorexigenic inflammatory cytokines, yet how it engages neural circuits that suppress feeding remains incompletely understood. Specifically, the role of peripheral sensory neurons in mediating IL-1β-induced anorexia is unresolved. Here, using DREADD-mediated inhibition of discrete peripheral sensory neuron populations and fiber photometry, we show that IL-1β-induced anorexia occurs in at least two temporally and mechanistically distinct phases. Shortly after IL-1β administration, prostaglandin signaling through non-vagal sensory afferents rapidly inhibits hypothalamic AgRP neurons to suppress food intake. At later time points, anorexia becomes partially prostaglandin-independent and vagal afferent neuron-dependent. Our findings demonstrate that multiple molecular signals mediate IL-1β-induced anorexia, and that diverse peripheral sensory pathways, including a previously unappreciated contribution from non-vagal afferents, are critical links between systemic inflammation and neural control of appetite.

## INTRODUCTION

Inflammation elicits a conserved suite of sickness behaviors, including loss of appetite, lethargy, and fever ^1^. Although these responses may be adaptive acutely ^2–4^, in chronic disease contexts they are maladaptive. This is particularly true of inflammation-induced anorexia, where appetite loss and the resulting malnutrition and sarcopenia may worsen outcomes in a variety of disease states ^4–7^. Inflammatory signals, including cytokines and prostaglandins released in the periphery, mediate anorexia ^8,9^. However, despite decades of study, the neural pathways by which anorectic inflammatory signals alter feeding behavior remain incompletely defined.

Interleukin-1β (IL-1β) is one of the most potent anorectic cytokines and suppresses feeding after intracranial ^10–14^, intravenous ^15,16^, or intraperitoneal ^14,16–20^ administration. Signaling in circumventricular regions is a commonly proposed mechanism for IL-1β-induced anorexia given the similar behavioral effects of different administration routes and the proximity of circumventricular regions to feeding circuitry in the hypothalamus and brainstem ^8,12,15,21,22^.

Peripheral sensory neurons, particularly vagal afferent neurons, have also been implicated in inflammation-induced anorexia and other sickness behaviors ^23^. Ex vivo experiments and studies in anesthetized animals consistently show that peripherally administered IL-1β and other inflammatory signals can activate vagal afferent neurons ^6,24–29^. However, studies in vagotomized rodents have yielded mixed results with regard to inflammation-induced anorexia ^7,30–36^, and interpretation of these studies is difficult due to loss of efferent in addition to afferent vagal signaling ^23^. Thus, whether vagal or other peripheral afferent activity is necessary for IL-1β-induced anorexia remains unclear.

In addition to the neural circuits being largely unknown, the molecular signals downstream of IL-1β that cause anorexia are incompletely understood. Multiple studies have shown that pharmacological inhibition or genetic deletion of cyclooxygenase enzymes, or of downstream prostaglandin-synthesis enzymes, significantly blunts IL-1β-induced anorexia, possibly via action at the blood-brain barrier ^9,15,16,19,20,37–40^. However, recent work indicates that prostaglandins also act on sensory neurons to drive anorexia. Specifically, during influenza infection, airway-innervating Gabra1-positive sensory afferents in the nodose-jugular-petrosal (NJP) ganglion complex mediate anorexia in mice ^6^. Thus, both IL-1β and prostaglandins may suppress feeding through multiple neural circuits and molecular mechanisms.

Here, we sought to define the peripheral circuits engaged by inflammatory signals *in vivo*. Because the activity of AgRP neurons in the arcuate nucleus of the hypothalamus closely tracks nutritional state and food intake ^41–47^, we began by using fiber photometry in this population as a readout of circuit-level changes in response to peripherally administered inflammatory stimuli.

We found that IL-1β inhibits AgRP neurons and causes rapid feeding suppression via a prostaglandin-dependent mechanism, and that this early anorexia is largely mediated by non-vagal peripheral sensory neurons. By contrast, vagal afferent neurons may play a role in the more prolonged effects of IL-1β on food intake. Taken together, our work highlights the complex and distributed mechanisms through which inflammation modulates behavior and offers insights into the circuits affected most acutely.

## RESULTS

### IL-1β causes severe anorexia and inhibits AgRP neurons

To dissect the mechanisms through which IL-1β causes anorexia, we first confirmed that IL-1β suppressed food intake in overnight-fasted mice (**Figure 1A–1C**). Relatively few studies have directly examined the effects of inflammatory signals on AgRP neuron activity ^6,21,22^, and the bacterial cell wall component lipopolysaccharide (LPS) appears to reduce food intake through mechanisms independent of AgRP neuron activity ^42,44,48,49^. Using fiber photometry, we found that intraperitoneal IL-1β administration significantly inhibited AgRP neurons with timing that aligned with anorexia onset (**Figure 1D–1F**). Although IL-1β-induced anorexia persisted for more than two hours, AgRP activity returned to baseline within 45 minutes of injection (**Figure 1D–1F**), suggesting that additional neuron populations are involved in this process.

**Figure 1.**
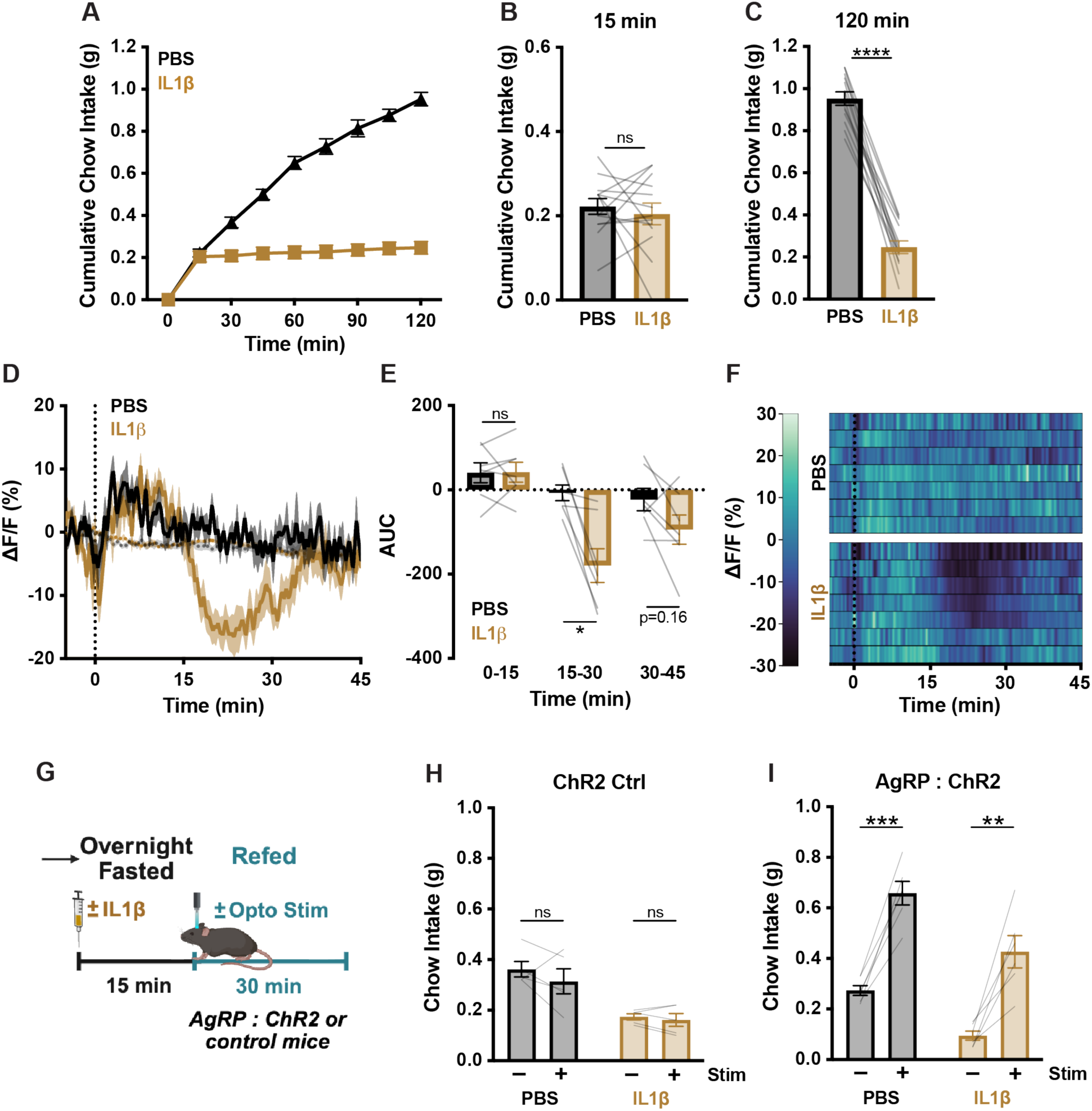
IL-1β reduces food intake in part via AgRP neuron inhibition. **(A–C)** Cumulative chow intake at the indicated time points in fasted C57BL/6 mice following i.p. injection with PBS or IL-1β (5 μg/kg) at time 0. Two-way repeated-measures ANOVA: main effects of time, treatment, and time×treatment interaction, p < 0.0001. n = 13 mice, paired design. **(D)** AgRP neuron calcium signal in fasted mice injected i.p. with PBS or IL-1β (5 μg/kg) at time 0. Isosbestic traces are shown in gray, vertical dashed line indicates injection time, and traces represent mean ± SEM. n = 7 mice, paired design. **(E)** Area under the curve (AUC) of traces from **(D)** across 3 15-min time windows following injection. Two-way repeated-measures ANOVA: main effect of time window, p < 0.0001; main effect of treatment, p= 0.0335; time×treatment interaction, p = 0.0011. **(F)** Heatmaps of ΔF/F (%) in individual mice from **(D)**, with subject order preserved across conditions. **(G)** Optogenetic experiment schematic **(H, I)** Thirty-minute chow intake in fasted ChR2 control mice (that do not express Cre recombinase) **(H)** and AgRP:ChR2 mice **(I)** following i.p. injection with PBS or IL1β (5 μg/kg) with or without light stimulation. **(H)** Two-way repeated-measures ANOVA: main effect of treatment, p = 0.0053; main effect of stimulation, p = 0.24; stimulation×treatment interaction, p = 0.53. n = 5 mice, paired design. **(I)** Two-way repeated-measures ANOVA: main effect of stimulation, p = 0.0001; main effect of treatment, p = 0.0099; stimulation x treatment interaction, p = 0.56. n = 6 mice, paired design. **(A, B, C, E, H, I)** Error bars indicate mean ± SEM. Lines represent individual mice. Holm–Šídák post hoc comparisons: *p<0.05, **p<0.01, ***p<0.001, ****p<0.0001

### AgRP neuron stimulation partially rescues IL-1β-induced anorexia

Activation of AgRP neurons induces vigorous feeding ^50,51^. It also rescues anorexia induced by stimuli, such as cholecystokinin (CCK) and glucagon-like peptide-1 (GLP-1) receptor agonists, which are sufficient to inhibit AgRP neurons, but does not rescue lipopolysaccharide (LPS)-induced anorexia, which does not directly suppress AgRP activity *in vivo* ^42,44,52^. Whether AgRP neuron stimulation can counteract anorexia caused by other inflammatory signals has not been examined. To test this, we used an optogenetic approach and showed that AgRP neuron stimulation partially rescues food intake in IL-1β-treated mice (**Figure 1G–1I**). This suggests that AgRP neuron inhibition may partially mediate IL-1β-induced anorexia.

### Peripheral sensory neuron inhibition rescues early IL-1β-induced anorexia

Systemic IL-1β induces profound anorexia and inhibits AgRP neurons, but the circuits through which this occurs remain unclear. Nutrient-related signals modulate food intake and AgRP neuron activity through peripheral sensory neurons, including both vagal and spinal afferent neurons ^53–55^. Whether IL-1β-induced activation of vagal or other peripheral afferent neurons is required to induce anorexia is unclear ^7,30–36,56^.

To test the contribution of a broad population of spinal and vagal sensory afferent neurons to IL-1β-induced anorexia, we generated mice expressing the inhibitory DREADD receptor (hM4Di) in NaV1.8-expressing neurons (NaV1.8:hM4Di mice) ^57,58^. The chemogenetic actuator deschloroclozapine (DCZ) alone had no effect on fast re-feeding in either NaV1.8:hM4Di mice or control NaV1.8-Cre-expressing mice lacking hM4Di (NaV1.8:Ctrl) (**Figure S1A–S1D**) ^59^. To confirm that NaV1.8 sensory neuron inhibition was effective, we showed that DCZ pretreatment significantly reduced cholecystokinin (CCK)-induced anorexia in NaV1.8:hM4Di but not NaV1.8:Ctrl mice (**Figure S1E–S1H**), consistent with prior findings that CCK-induced anorexia requires vagal afferent signaling ^32,55,60^.

Next, we examined the role of NaV1.8 neurons in IL-1β-induced anorexia. Because the onset of IL-1β-induced anorexia occurs approximately 15 minutes after injection (Figure 1A), DCZ was administered 20 minutes before IL-1β and mice were re-fed 15 minutes later. DCZ pretreatment almost completely restored food intake for approximately 30 minutes in NaV1.8:hM4Di mice with minimal effects on feeding at later time points (**Figure 2D–2F**), whereas NaV1.8:Ctrl mice showed no change in IL-1β-induced anorexia (**Figure 2A–2C**). Thus, peripheral sensory neuron signaling is selectively required for early IL-1β-induced anorexia.

**Figure 2.**
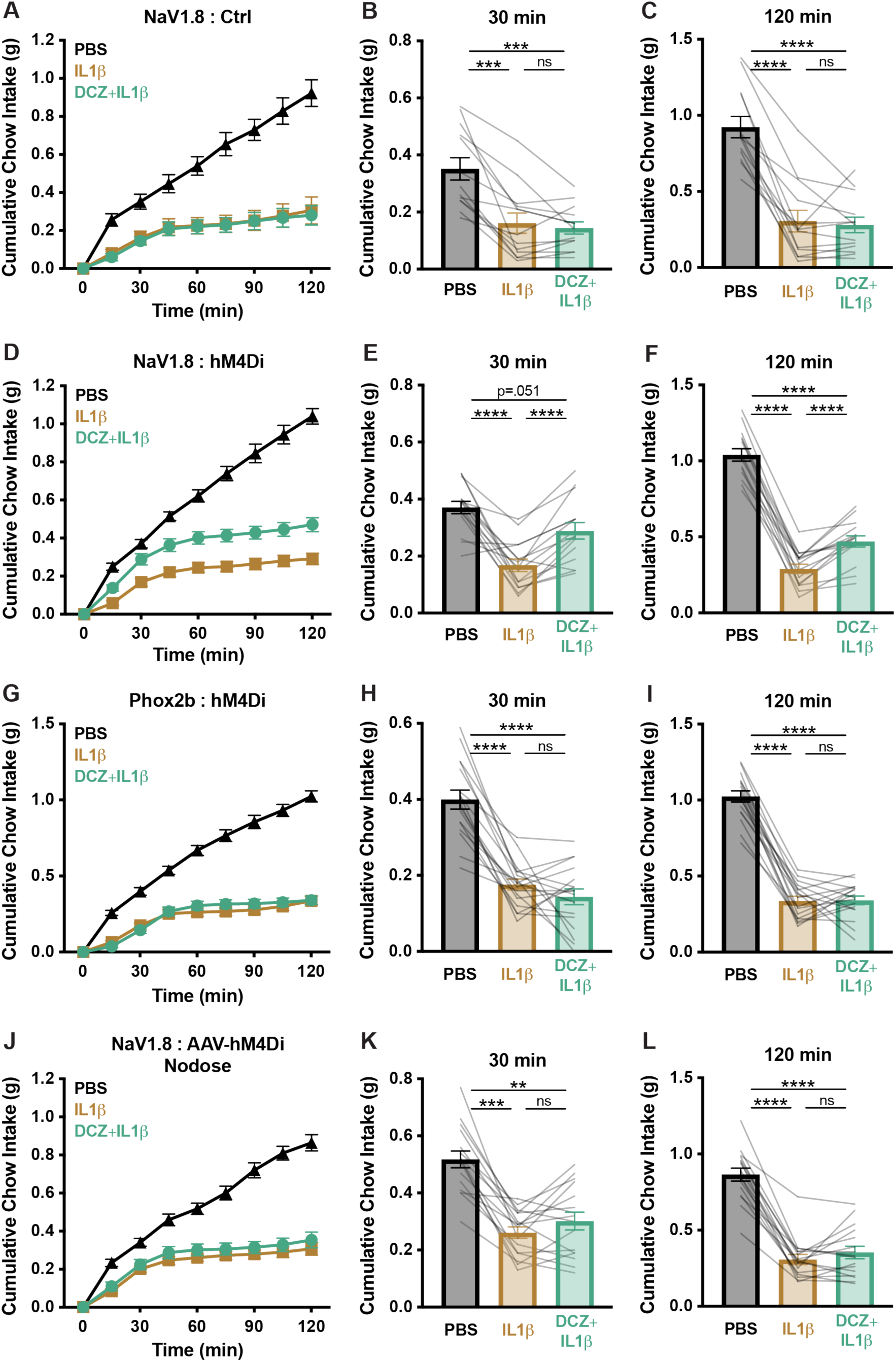
Early IL-1β-induced feeding suppression is primarily mediated by non-vagal NaV1.8-expressing peripheral sensory neurons. **(A–F)** Cumulative chow intake at the indicated time points in fasted NaV1.8:Ctrl **(A–C)** and NaV1.8:hM4Di **(D–F)** mice following i.p. injection with PBS, IL-1β (5 μg/kg), or DCZ (100 μg/kg)+IL-1β. **(A)** Two-way repeated-measures ANOVA: main effects of time, treatment, and time×treatment interaction, p < 0.0001. n = 13 mice, paired design. **(D)** Two-way repeated-measures ANOVA: main effects of time, treatment, and time×treatment interaction, p < 0.0001. n = 15 mice, paired design. **(G–I)** Cumulative chow intake at the indicated time points in fasted Phox2b:hM4Di mice following i.p. injection of PBS, IL-1β (5 μg/kg), or DCZ (100 μg/kg)+IL-1β. **(G)** Two-way repeated-measures ANOVA: main effects of time, treatment, and time×treatment interaction, p < 0.0001. n = 17 mice, paired design. **(J–L)** Cumulative chow intake at the indicated time points in fasted NaV1.8:AAV-hM4Di nodose mice following i.p. injection of PBS, IL-1β (5 μg/kg), or DCZ (100 μg/kg)+IL-1β. Two-way repeated-measures ANOVA: main effects of time, treatment, and time×treatment (all p < 0.0001). n = 16 mice, paired design. **(A–L)** Error bars indicate mean ± SEM. Lines represent individual mice. Holm–Šídák post hoc comparisons: **p<0.01, ***p<0.001, ****p<0.0001

### Early IL-1β-induced anorexia requires the activity of non-vagal sensory neurons

To determine whether the effect of sensory neuron inhibition on IL-1β-induced anorexia occurs through vagal or spinal afferents, we generated mice expressing hM4Di in Phox2b-expressing neurons (Phox2b:hM4Di) ^61,62^. Phox2b is expressed throughout the nodose-petrosal ganglion complex, labeling all visceral vagal afferents, including NaV1.8-expressing subdiaphragmatic nodose neurons, but it is not expressed in NaV1.8-expressing sensory neurons of the jugular ganglion ^63,64^ or in afferents outside the vagal ganglia, such as those in the dorsal root ganglia (DRG) ^55^.

DCZ alone had no effect on fast re-feeding in either Phox2b:hM4Di or Phox2b:Ctrl mice that express Phox2b-Cre but lack hM4Di expression (**Figure S2A–S2D**). As with NaV1.8:hM4Di mice, and as expected, DCZ pretreatment partially ameliorated CCK-induced anorexia in Phox2b:hM4Di mice but not in Phox2b:Ctrl mice (**Figures S2E–S2H**). Chemogenetic inhibition of Phox2b-expressing neurons failed to rescue IL-1β-induced anorexia, and DCZ had no effect on anorexia in Phox2b:Ctrl mice (**Figures 2G–2I; S2I–S2K**).

Because Phox2b is expressed outside the nodose and petrosal ganglia, including in the geniculate ganglia and brainstem regions that regulate feeding ^61,62^, and NaV1.8 is found outside of the NJP ganglia in other sensory afferent populations^65^, we next surgically restricted chemogenetic manipulation to the NJP ganglia. Briefly, mice expressing NaV1.8-Cre underwent bilateral nodose ganglia injections with AAVs encoding either Cre-dependent hM4Di (NaV1.8:AAV-hM4Di) or EGFP (NaV1.8:AAV-Ctrl) ^66^. RT-qPCR was used to verify transgene expression. Human CHRM4 expression (a component of the hM4Di transgene) was detected in all AAV-hM4Di-injected vagal ganglia but was undetectable in any AAV-Ctrl controls, whereas expression of the NaV1.8-encoding gene *Scn10a* was detected in all NJP ganglia samples (**Figure S3B, S3C**).

DCZ had no effect on fast re-feeding in NaV1.8:AAV-Ctrl mice (**Figure S3D**). By contrast, unlike in Phox2b:hM4Di mice, DCZ slightly but significantly increased re-feeding in NaV1.8:AAV-hM4Di animals (**Figure S3E**). As with the Phox2b experiments, DCZ pretreatment partially rescued CCK-induced anorexia in NaV1.8:AAV-hM4Di but not in NaV1.8:AAV-Ctrl mice (**Figure S3F–S3K**), confirming effective inhibition of nodose neurons. DCZ had no effect on IL-1β-induced anorexia in either NaV1.8:AAV-hM4Di or NaV1.8:AAV-Ctrl mice (**Figures 2J–2L; S3L–S3N**). Thus, using both transgenic and virus-mediated chemogenetic inhibition of vagal afferent neurons, we showed that sensory neuron mediation of early IL-1β-induced anorexia is at least predominantly non-vagal.

### Prostaglandins mediate early IL-1β-induced anorexia and IL-1β-induced AgRP neuron inhibition

Our findings that IL-1β administration transiently inhibits AgRP neurons (**Figure 1D–1F**) and that non-vagal sensory neuron inhibition selectively rescues early IL-1β-induced anorexia (**Figure 2**) support multiple mechanisms of IL-1β-induced anorexia over time.

However, the specific molecular mediators of early and late IL-1β-induced anorexia are unknown. Prior studies suggest that prostaglandins are critical for IL-1β-induced sickness behaviors, including anorexia ^9,16,19,38,67,68^. We therefore tested how prostaglandin signaling contributes to IL-1β-mediated anorexia and AgRP neuron inhibition.

To determine whether prostaglandins are required for IL-1β-induced anorexia, we pretreated fasted wild-type mice with ketorolac (a COX-1/2 inhibitor) 20 min before IL-1β. Re-feeding began 15 min after IL-1β administration. Ketorolac pretreatment completely restored food intake for 30 minutes after re-feeding, after which intake declined compared to PBS-treated mice (**Figure 3A–3C**). These results indicate an early prostaglandin-dependent phase of anorexia followed by a later phase with partial prostaglandin dependence, in agreement with prior studies ^37^. Pretreatment with the COX-2-selective inhibitor parecoxib produced partial restoration of re-feeding similar to ketorolac (**Figure S4G–S4I**).

**Figure 3.**
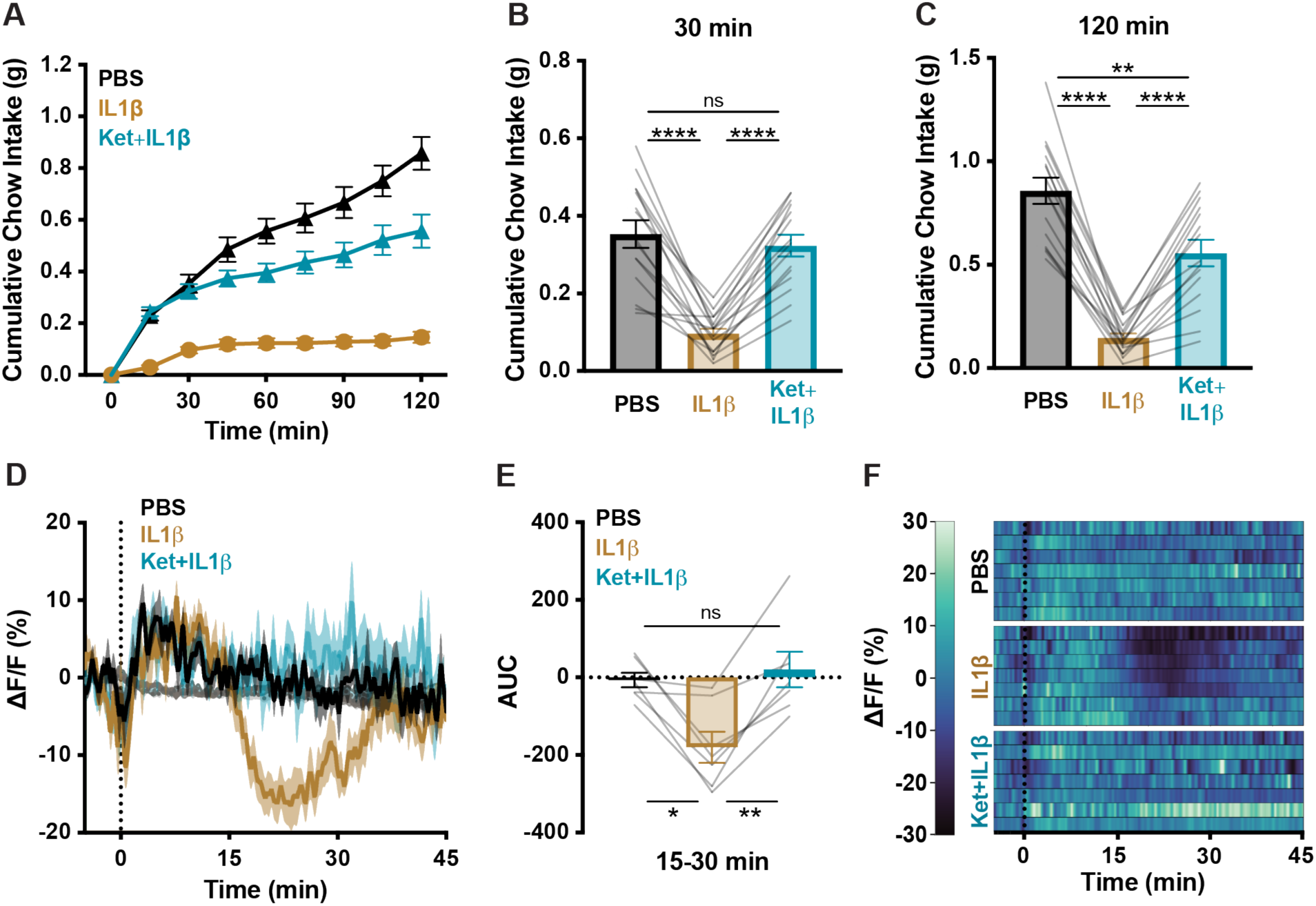
Prostaglandin release is necessary for IL-1β-mediated AgRP neuron inhibition and early feeding suppression. **(A–C)** Cumulative chow intake at the indicated time points in fasted C57BL/6 mice following i.p. injection with PBS, IL-1β (5 μg/kg), or ketorolac (Ket, 15 mg/kg)+IL-1β. Two-way repeated-measures ANOVA: main effects of time, treatment, and time×treatment, p < 0.0001. n = 15 mice, paired design. **(D)** AgRP neuron calcium signal in fasted mice injected i.p. with PBS, IL-1β (5 μg/kg), or ketorolac (15 mg/kg)+IL-1β at time 0. Isosbestic traces are shown in gray, vertical dashed line indicates injection time, and traces represent mean ± SEM. n = 7 mice, paired design. **(E)** Area under the curve (AUC) of traces from **(D)** at the 15 to 30-min time window following injection. Two-way repeated-measures ANOVA across the 0–15, 15–30, and 30–45 min time windows: main effect of time, p = 0.0061; main effect of treatment, p = 0.0484; time×treatment interaction, p = 0.0005. **(F)** Heatmaps of ΔF/F (%) in individual mice from **(D)**, with subject order preserved across conditions. **(A, B, C, E)** Error bars indicate mean ± SEM. Lines represent individual mice. Holm–Šídák post hoc comparisons: *p<0.05, **p<0.01, ****p<0.0001

We next tested whether IL-1β-mediated AgRP neuron inhibition also requires prostaglandin signaling. Strikingly, ketorolac pretreatment completely prevented IL-1β-induced AgRP neuron inhibition (**Figure 3D–3F**). Ketorolac alone had no effect on AgRP neuron activity or food intake (**Figure S4A–S4F**). Taken together, these findings indicate that prostaglandin signaling is necessary for both early IL-1β-induced anorexia and IL-1β-driven AgRP neuron inhibition.

### Multiple prostaglandins cause anorexia and inhibit AgRP neurons

PGE2 has been the primary prostaglandin examined in inflammation-induced anorexia and other sickness behaviors, though additional prostaglandins have been implicated in IL-1β-induced anorexia ^38,69^ and their effects may be strain-dependent ^38^. To better understand which prostaglandins might contribute to IL-1β-induced anorexia, we tested the anorectic effects of four prostaglandins (PGE2, PGD2, PGF2α, and PGI2) and examined their effect on AgRP neuron activity. All four prostaglandins significantly decreased food intake (**Figure 4A–4D**) and inhibited AgRP neurons (**Figures 4E–4L**).

**Figure 4.**
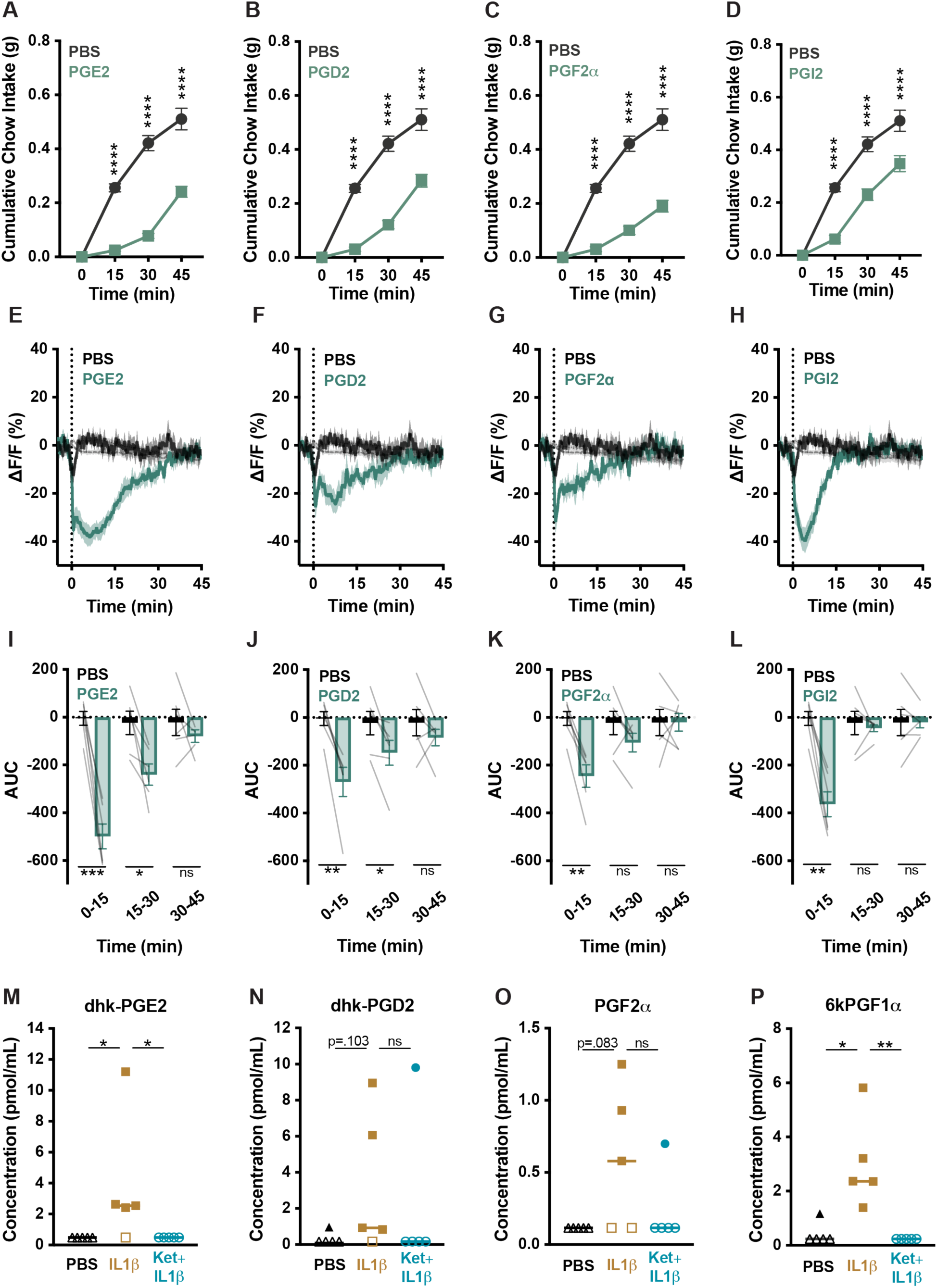
IL-1β induces prostaglandin release, and prostaglandins suppress food intake and inhibit AgRP neurons. **(A–D)** Cumulative chow intake at the indicated time points in fasted C57BL/6 mice following i.p. injection with PBS or prostaglandins: PGE2 (250 μg/kg) **(A)**; PGD2 (250 μg/kg) **(B)**; PGF2α (1 mg/kg) **(C)**; PGI2 (250 μg/kg) **(D)**. Two-way repeated-measures ANOVA: main effects of treatment and time, p < 0.0001 for all panels; treatment×time interaction, p = 0.0008 **(A)**, p = 0.0093 **(B)**, p = 0.0020 **(C)**, p = 0.0044 **(D)**. n = 15 mice per group, paired design. Holm–Šídák post hoc comparisons with significance shown above each time point. **(E–H)** AgRP neuron calcium signal in fasted mice injected i.p. with PBS or prostaglandins at time 0: PGE2 (250 μg/kg) **(E)**; PGD2 (250 μg/kg) **(F)**; PGF2α (1 mg/kg) **(G)**; PGI2 (250 μg/kg) **(H)**. Isosbestic traces are shown in gray, vertical dashed lines indicate injection time, and traces represent mean ± SEM. n = 6 mice per group, paired design. **(I–L)** Area under the curve (AUC) of traces from **(E–H)** across 3 15-min time windows following injection. **(I)** Two-way repeated-measures ANOVA for PGE2: main effect of time window, p = 0.0027; main effect of treatment, p = 0.0013; time×treatment interaction, p = 0.0022. **(J)** Two-way repeated-measures ANOVA for PGD2: main effect of time window, p = 0.054; main effect of treatment, p = 0.012; time×treatment interaction, p = 0.0056. **(K)** Two-way repeated-measures ANOVA for PGF2α: main effect of time window, p = 0.047; main effect of treatment, p = 0.056; time×treatment interaction, p = 0.0022. **(L)** Two-way repeated-measures ANOVA for PGI2: main effect of time window, p = 0.0003; main effect of treatment, p = 0.0027; time×treatment interaction, p = 0.028. **(M–P)** Plasma concentrations of dhk-PGE2 (a stable metabolite of PGE2; **(M)**), dhk-PGD2 (a stable metabolite of PGD2; **(N)**), PGF2α **(O)**, and 6k-PGF1α (a stable metabolite of PGI2; **(P)**) following i.p. injection with PBS, IL-1β (5 μg/kg), or ketorolac (Ket, 15 mg/kg) + IL-1β. Detection frequencies compared by one-tailed Fisher’s exact tests (PBS vs. IL-1β and IL-1β vs. ketorolac + IL-1β) with Holm–Šídák correction: **(M)** dhk-PGE2, PBS 0/5, IL-1β 4/5, ketorolac + IL-1β 0/5; **(N)** dhk-PGD2, PBS 1/5, IL-1β 4/5, ketorolac + IL-1β 1/5; **(O)** PGF2α, PBS 0/5, IL-1β 3/5, ketorolac + IL-1β 1/5; **(P)** 6k-PGF1α, PBS 1/5, IL-1β 5/5, ketorolac + IL-1β 0/5. Each symbol represents an individual mouse; filled symbols indicate detectable values and hollow symbols indicate non-detectable values (below the limit of detection), and horizontal bars denote medians. Detection-frequency statistics were computed on detection status (detectable vs non-detectable), while imputed values were only used for plotting and summary statistics. Non-detectable values were imputed as one-fifth of the minimum detected concentration across all samples, following UCSD Lipidomics Core guidelines. n = 5 mice per condition. **(A–D, I–L)** Error bars indicate mean ± SEM. Lines represent individual mice. Holm–Šídák post hoc comparisons: *p<0.05, **p<0.01, ***p<0.001, ****p<0.0001

To determine which of these prostaglandins could be responsible for IL-1β-induced anorexia, we performed a plasma lipidomic analysis collected 30 minutes after IL-1β injection. Multiple eicosanoid species were significantly upregulated, including stable metabolites of PGE2 (dhk-PGE2; **Figure 4M**; ^70,71^) and PGI2 (6kPGF1α; **Figure 4P**; ^70,71^), while increases in PGD2 (dhk-PGD2; **Figure 4N**; ^71^) and PGF2α (**Figure 4O**; ^71^) did not reach significance. In contrast, plasma from PBS-injected mice contained minimal or undetectable levels of all these metabolites.

Pretreatment with ketorolac eliminated the IL-1β-induced increases in prostaglandin levels, as expected (**Figures 4M–4P**). Taken together, these data suggest that multiple prostaglandins may serve as molecular mediators of IL-1β-induced anorexia.

### Peripheral sensory neurons contribute to prostaglandin-induced anorexia

Since prostaglandins and peripheral sensory neuron activity are required for early IL-1β-induced anorexia, and both vagal and spinal afferents express a variety of prostaglandin receptors ^6,64,72,73^, we next tested whether sensory neurons mediate the anorectic effects of prostaglandins. Chemogenetic inhibition of NaV1.8-expressing sensory neurons in NaV1.8:hM4Di mice completely prevented PGE2- and PGD2-induced anorexia (**Figure 5A, 5B, 5E, 5F**). In contrast, NaV1.8 neuron inhibition only partially rescued PGI2- and PGF2α-induced anorexia (**Figure 5C, 5D, 5G, 5H**), suggesting that different prostaglandins suppress food intake through distinct pathways. DCZ administration did not affect prostaglandin-induced anorexia in NaV1.8:Ctrl mice (**Figure S5A–S5H**).

**Figure 5.**
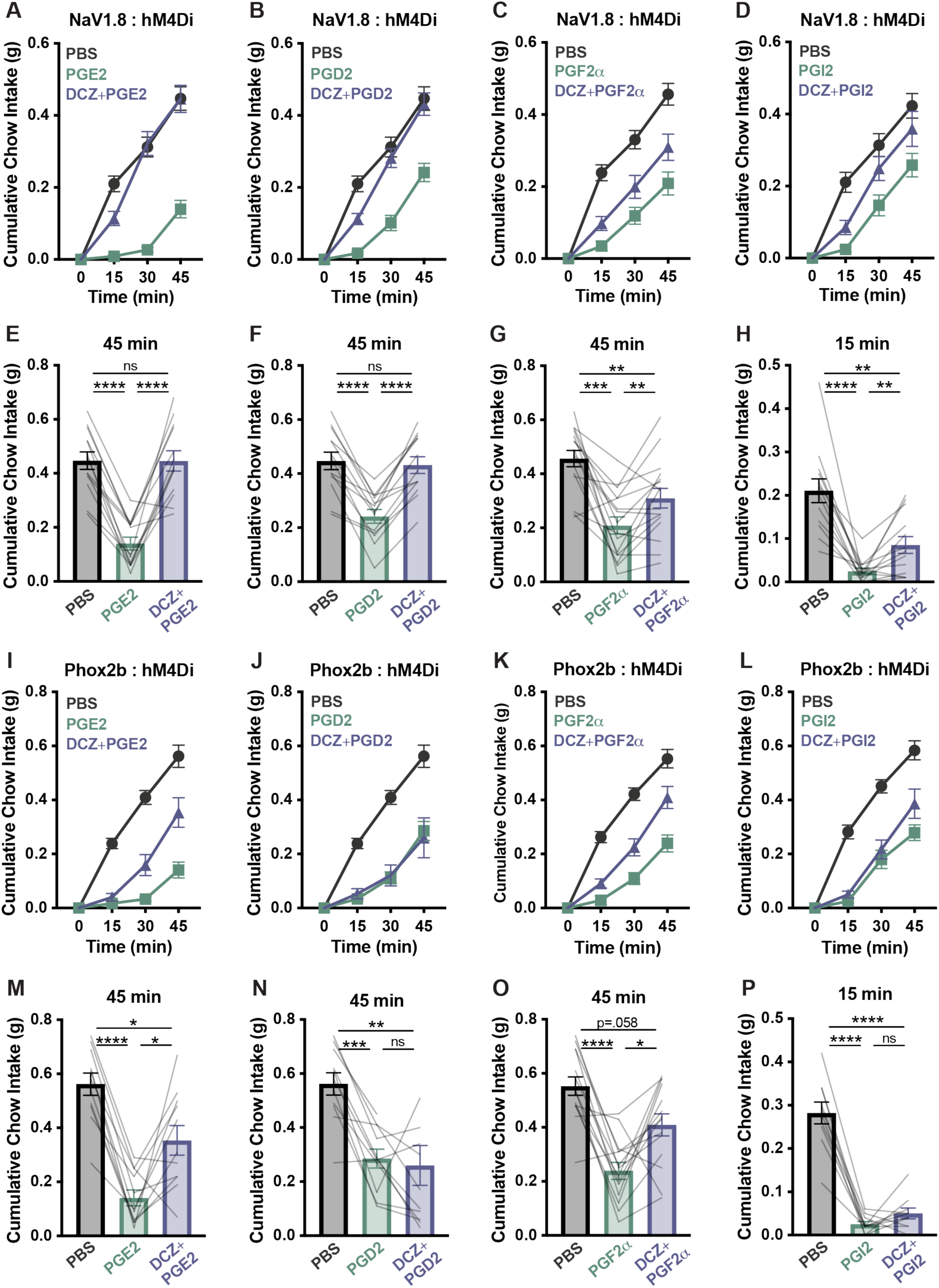
Multiple prostaglandins induce anorexia via NaV1.8-expressing peripheral sensory neurons, with prostaglandin-specific vagal contributions. **(A–H)** Cumulative chow intake at the indicated time points in fasted NaV1.8:hM4Di mice following i.p. injection with PBS, PGE2 (250 μg/kg) **(A, E)**, PGD2 (250 μg/kg) **(B, F)**, PGF2α (1 mg/kg) **(C, G)**, or PGI2 (250 μg/kg) **(D, H)**, with or without DCZ (100 μg/kg). **(A–C)** Two-way repeated-measures ANOVA: main effects of time, treatment, and time×treatment interaction, p < 0.0001. **(D)** Two-way repeated-measures ANOVA: main effect of time, p < 0.0001; main effect of treatment, p = 0.0012; time×treatment interaction, p = 0.0025. n = 13-16 mice, paired design. **(I–P)** Cumulative chow intake at the indicated time points in fasted Phox2b:hM4Di mice following i.p. injection with PBS, PGE2 (250 μg/kg) **(I, M)**, PGD2 (250 μg/kg) **(J, N)**, PGF2α (1 mg/kg) **(K, O)**, or PGI2 (250 μg/kg) **(L, P)**, with or without DCZ (100 μg/kg) pretreatment. **(I–L)** Two-way repeated-measures ANOVA: main effects of time, treatment, and time×treatment interaction, p < 0.0001. n = 10-14 mice, paired design. **(A–P)** Error bars indicate mean ± SEM. Lines represent individual mice. Holm–Šídák post hoc comparisons: *p<0.05, **p<0.01, ***p<0.001, ****p<0.0001.

Although vagal afferent neurons did not appear to play a major role in the early prostaglandin-mediated phase of IL-1β-induced anorexia (**Figure 2G–2L**), prior work reported that NJP ganglion complex sensory neurons drive PGE2-induced anorexia during influenza infection^6^. Because the petrosal and nodose ganglia are closely related epibranchial sensory ganglia that both express Phox2b and share visceral sensory functions^64^, we examined whether Phox2b-expressing neurons might contribute to prostaglandin-induced anorexia. Chemogenetic inhibition of Phox2b neurons partially ameliorated PGE2- and PGF2α-induced anorexia (**Figure 5I, 5K, 5M, 5O**) but did not rescue PGD2- or PGI2-induced anorexia (**Figure 5J, 5L, 5N, 5P**).

DCZ treatment did not alter prostaglandin-induced anorexia in Phox2b:Ctrl mice (**Figure S5I–S5P**). Phox2b neuron inhibition produced a smaller reduction in PGE2-induced anorexia than NaV1.8 neuron inhibition, suggesting that vagal afferent neurons only partially account for PGE2-induced anorexia. To test this possibility more directly, we performed nodose-targeted viral inhibition experiments in NaV1.8-Cre mice using AAV-hM4Di or AAV-Ctrl. Chemogenetic inhibition of nodose neurons in NaV1.8:AAV-hM4Di mice partially attenuated PGE2-induced anorexia (**Figure S6A–S6D**), while PGD2-induced anorexia was unaffected (**Figure S6E–S6H**), consistent with our results in Phox2b:hM4Di animals (**Figure 5I, 5J, 5M, 5N**). The food intake rescue in nodose ganglia-injected mice was less robust than in Phox2b:hM4Di mice. This may be due to more widespread hM4Di expression in the transgenic model, or due to the fact that the *Gabra1*-expressing petrosal-nodose sensory neurons that have been shown to mediate PGE2-induced anorexia largely do not express NaV1.8 ^63^

### The prostaglandin-independent component of IL-1β-induced anorexia is not mediated by direct IL-1β signaling on NaV1.8 sensory neurons

The previous experiments suggest that the onset of IL-1β-induced anorexia is prostaglandin dependent and requires signaling through largely non-vagal NaV1.8 sensory neurons. IL-1β-induced anorexia at later time points is only partially prostaglandin-dependent, as indicated by incomplete rescue by ketorolac and parecoxib after the initial 30 minutes of re-feeding (**Figures 3A–3C; S4G–S4I**). We thus set out to investigate the later, partially prostaglandin-independent component of IL-1β-induced anorexia.

Some studies suggest that subsets of NaV1.8-positive nociceptors may respond directly to IL-1β through IL-1 receptor (*Il1r1*) signaling^74,75^, and several reports describe IL-1β-evoked activation of vagal afferent neurons along with transcriptional detection of *Il1r1*^25–27^. However, other studies have not detected *Il1r1* in sensory neurons ^76,77^. We tested the hypothesis that prostaglandin-independent IL-1β-induced anorexia involves direct IL-1β signaling on sensory neurons by generating mice with *Il1r1* selectively knocked out of NaV1.8-expressing neurons (NaV1.8:IL1R1-cKO) and littermate controls lacking the floxed *Il1r1* allele (NaV1.8:IL1R1-Ctrl) and monitored feeding in these animals after IL-1β injection^78^. Mice were also pretreated with ketorolac to block prostaglandin synthesis and isolate the prostaglandin-independent contribution to anorexia. IL-1β-induced feeding suppression was similar in NaV1.8:IL1R1-cKO and NaV1.8:IL1R1-Ctrl mice (**Figure 6**), indicating that direct IL-1β signaling in NaV1.8 sensory neurons is not required for the prostaglandin-independent component of IL-1β-induced anorexia.

**Figure 6.**
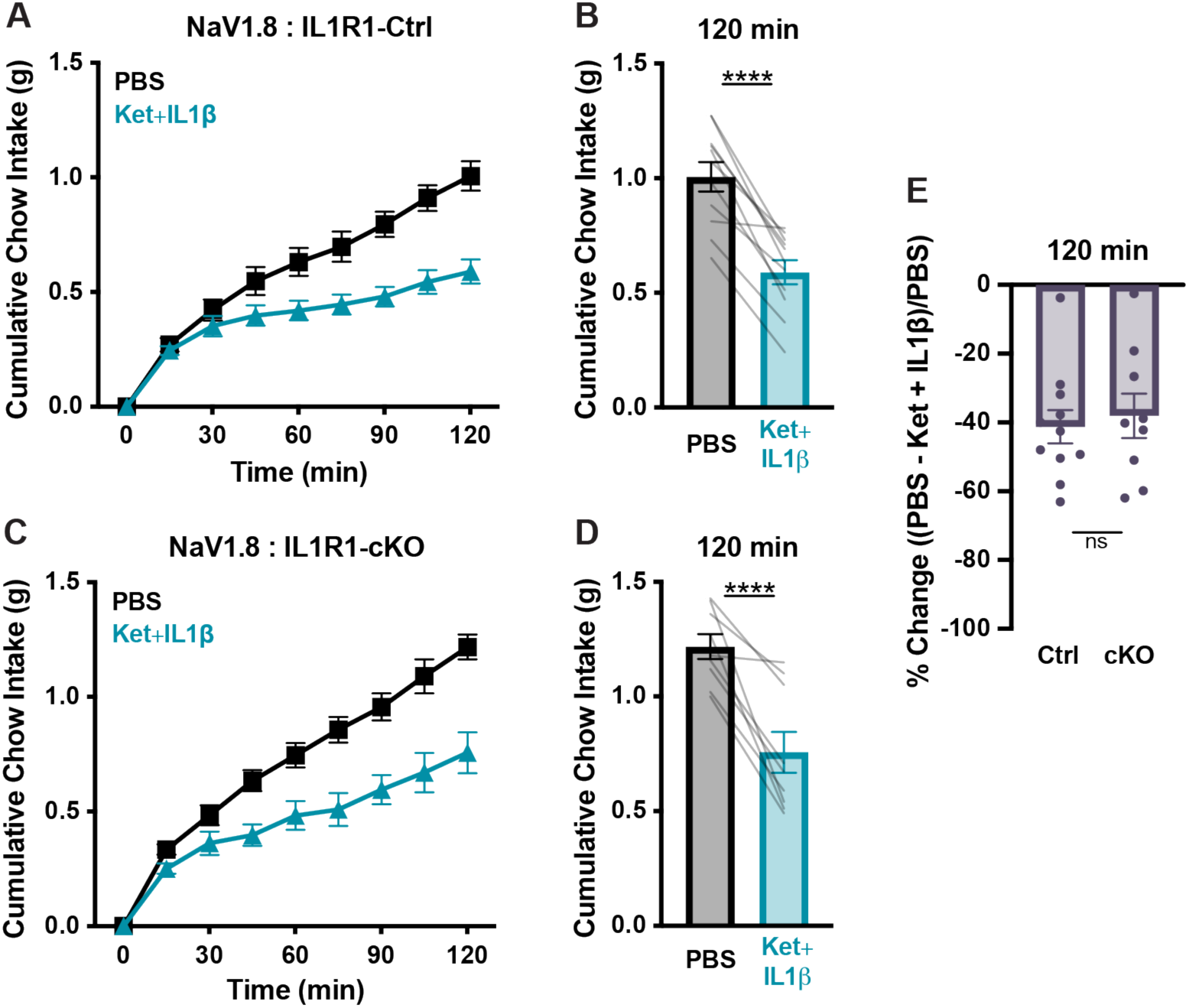
IL1R1 expression in peripheral sensory neurons is not required for prostaglandin-independent, IL-1β-induced anorexia. **(A–D)** Cumulative chow intake at the indicated time points in fasted NaV1.8:IL1R1-Ctrl **(A, B)** or NaV1.8:IL1R1-cKO **(C, D)** mice following i.p. injection with PBS or ketorolac (Ket, 15 mg/kg) +IL-1β (5 μg/kg). **(A)** Two-way repeated-measures ANOVA: main effect of time, p < 0.0001; main effect of treatment, p = 0.0002; time×treatment interaction, p < 0.0001. **(C)** Two-way repeated-measures ANOVA: main effect of time, p < 0.0001; main effect of treatment, p = 0.0002; time×treatment interaction, p = 0.0009. n = 11 control and 9 cKO mice. **(E)** Percent feeding suppression relative to PBS induced by ketorolac + IL-1β in mice from **(A–D)**. Unpaired two-tailed Welch’s t-test, p = 0.6966. **(A–E)** Error bars indicate mean ± SEM. Lines or dots represent individual mice. **(A, C)** Holm–Šídák post hoc comparisons: ****p<0.0001

### Vagal afferent activity contributes to prostaglandin-independent IL-1β-induced anorexia

Finally, we tested whether vagal afferents mediate the prostaglandin-independent component of IL-1β-induced anorexia through a mechanism other than direct IL-1β action on afferent neurons. We once again isolated the prostaglandin-independent anorexia component by pretreatment with ketorolac, and combined this with chemogenetic sensory neuron inhibition in our NaV1.8:hM4Di, Phox2b:hM4Di, and NaV1.8:AAV-hM4Di nodose mice. Fasted mice received either ketorolac plus DCZ or ketorolac alone prior to IL-1β injection followed by re-feeding. The addition of DCZ to ketorolac potentiated the rescue of anorexia in NaV1.8:hM4Di and Phox2b:hM4Di mice, suggesting that vagal afferent neurons play a prostaglandin-independent role in IL-1β-induced anorexia that we unmasked by blocking the prostaglandin-dependent effects (**Figure 7A–7G**). Enhanced food intake in NaV1.8:AAV-hM4Di mice pretreated with DCZ plus ketorolac versus ketorolac alone did not reach significance (**Figure 7H–7J**). In control mice not expressing hM4Di, DCZ had no effect on food intake (**Figure S7**). Taken together, these findings suggest that a substantial portion of the delayed, prostaglandin-independent component of IL-1β-induced anorexia is mediated by Phox2b-positive, NaV1.8-positive vagal afferents through a yet to be identified molecular mediator. By contrast, early, prostaglandin-dependent anorexia is mediated by non-vagal NaV1.8 sensory afferents and AgRP neuron inhibition.

**Figure 7.**
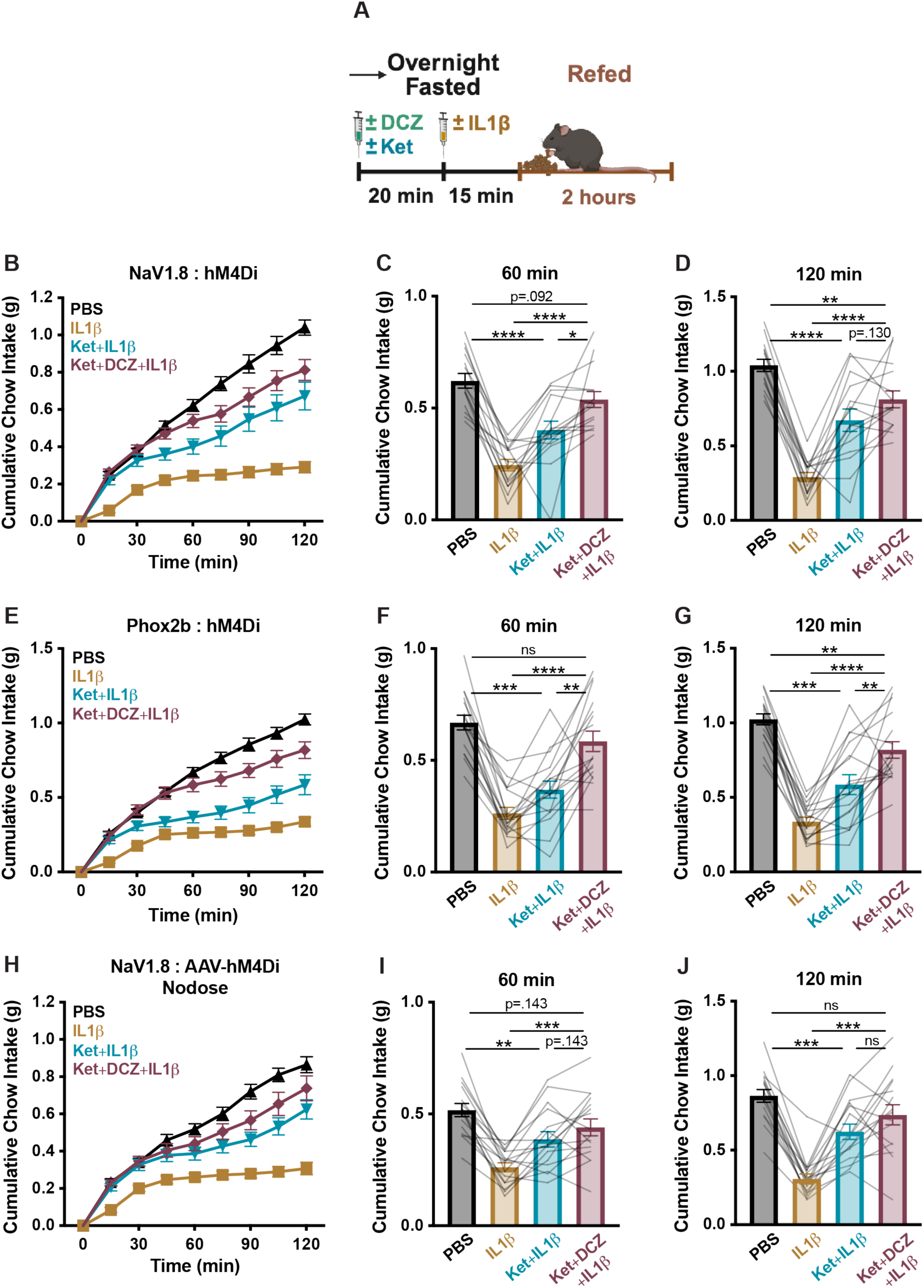
Inhibiting vagal afferent neurons rescues the prostaglandin-independent component of IL-1β-induced anorexia. **(A)** Experimental schematic for Figure 7 experiments. **(B–D)** Cumulative chow intake at the indicated time points in fasted NaV1.8:hM4Di mice following i.p. injection with PBS, IL-1β (5 μg/kg), ketorolac (Ket, 15 mg/kg)+IL-1β, or DCZ (100 μg/kg)+ketorolac+IL-1β. **(B)** Two-way repeated-measures ANOVA: main effects of time, treatment, and time×treatment interaction, p < 0.0001. n = 15 mice, paired design. **(E–G)** Cumulative chow intake at the indicated time points in fasted Phox2b:hM4Di mice following i.p. injection with PBS, IL-1β (5 μg/kg), ketorolac (15 mg/kg)+IL-1β, or DCZ (100 μg/kg)+ketorolac+IL1β. **(E)** Two-way repeated-measures ANOVA: main effects of time, treatment, and time×treatment interaction, p < 0.0001. n = 17 mice, paired design. **(H–J)** Cumulative chow intake at the indicated time points in fasted NaV1.8:AAV-hM4Di nodose mice following i.p. injection with PBS, IL-1β (5 μg/kg), ketorolac (15 mg/kg)+IL-1β, or DCZ (100 μg/kg)+ketorolac+IL-1β. **(H)** Two-way repeated-measures ANOVA: main effects of time, treatment, and time×treatment interaction, p < 0.0001. n = 16 mice, paired design. **(B–J)** Error bars indicate mean ± SEM. Lines represent individual mice. Holm–Šídák post hoc comparisons: *p<0.05, **p<0.01, ***p<0.001, ****p<0.0001

## DISCUSSION

Inflammation-induced anorexia and other sickness behaviors have often been attributed to inflammatory signaling at the blood-brain barrier (BBB) or circumventricular regions, though more recent studies have highlighted the importance of peripheral sensory pathways in appetite regulation and in immune responses ^6,8,21,24,53–55^ . However, the extent to which each of these mechanisms mediates inflammation-induced behavioral changes remains incompletely understood. We examined the role of peripheral sensory afferents in IL-1β-induced anorexia.

Using fiber photometry in AgRP neurons, chemogenetic inhibition of defined sensory neuron populations, and pharmacologic cyclooxygenase blockade, we identified multiple pathways contributing to IL-1β-induced anorexia, with peripheral sensory afferents playing a more complex role than previously thought. These findings provide a foundation for future therapeutic strategies for anorexia and cachexia that consider peripheral sensory circuits alongside BBB-centered mechanisms.

### Inflammatory signals inhibit AgRP neurons via a prostaglandin-dependent mechanism to suppress food intake

We showed that peripheral administration of IL-1β inhibits AgRP neurons in awake animals, and that the timing of this inhibition coincides with the onset of anorexia. Moreover, optogenetic stimulation of AgRP neurons partially rescues IL-1β -induced anorexia (**Figure 1G–1I**), suggesting that their inhibition is required for the early anorectic effect of acute inflammation.

Both the onset of anorexia and AgRP neuron inhibition are completely blocked by cyclooxygenase inhibition, demonstrating that this process is prostaglandin-dependent (**Figures 3; S4**), and administration of a variety of prostaglandins is also sufficient to inhibit AgRP neurons and suppress food intake (**Figure 4**). These findings complement recent studies showing that prostaglandin signaling mediates influenza-induced anorexia ^6^ and prior studies demonstrating the complex interplay between inflammatory pain, hunger, and food intake with AgRP neurons as a key regulator of this crosstalk ^79^.

### Non-vagal, NaV1.8-expressing sensory afferents mediate early, prostaglandin-dependent IL-1β-induced anorexia

NaV1.8 is expressed in large subsets of vagal sensory neurons primarily innervating subdiaphragmatic viscera, including those in the nodose ganglion and fused nodose-jugular-petrosal ganglion complex ^53,63,64^, in the majority of nociceptive dorsal root ganglion neurons as well as some non-nociceptive types, and across other cranial sensory ganglia ^57,65^.

Chemogenetic inhibition of NaV1.8 afferents restored the first 30 minutes of re-feeding following IL-1β administration, after which animals almost completely stopped eating for at least the next 90 minutes (**Figure 2A–2F**). By contrast, selectively chemogenetically inhibiting vagal afferent neurons via transgenic and viral approaches did not rescue IL-1β induced anorexia (**Figures 2G–2L; S2; S3**), consistent with a recent study showing that inhibition of Phox2b-expressing neurons does not rescue LPS-induced anorexia ^80^. Previous studies, which have variably implicated the vagus nerve in inflammation-induced anorexia, relied largely on vagotomies that sever both afferent and efferent pathways potentially confounding results. Our chemogenetic approach, which selectively targets afferent neurons, suggests that non-vagal NaV1.8 afferents mediate early IL-1β-induced anorexia.

As noted above, prostaglandin production is essential for this early behavioral effect. Ketorolac pretreatment completely restored food intake for the first 30 minutes of re-feeding following IL-1β administration and prevented IL-1β -induced AgRP neuron inhibition (**Figure 3**).

Chemogenetic inhibition of NaV1.8-expressing neurons largely rescued prostaglandin-induced anorexia, and this was only partially recapitulated by inhibition of vagal afferent neurons (**Figures 5; S5; S6**). Overall, these findings support a model in which IL-1β triggers peripheral prostaglandin release that activates non-vagal NaV1.8 sensory neurons to induce anorexia and inhibit AgRP neurons (**Figures 1–3**). These findings complement a recent study showing that PGE2 inhibits AgRP neurons and mediates influenza-induced sickness behaviors via activation of upper airway-innervating *Gabra1*-expressing petrosal ganglia neurons. Interestingly, Gabra1-expressing neurons negligibly overlap with NJP neurons expressing NaV1.8 (∼7.7%), whereas Phox2b and Gabra1 expression overlap is higher (60.1%) ^63^. This suggests that inflammation in different tissues may be communicated to the brain via distinct neural circuits and molecular mediators, which will require further study to elucidate. Moreover, neither of these findings preclude a role for BBB or circumventricular signaling in inflammation-induced anorexia as has been reported previously ^8,9,12,15,21,22^.

### Prostaglandins cause anorexia via multiple mechanisms

Further highlighting the complexity of inflammation-induced behavioral changes is that while all four prostaglandins tested (PGE2, PGD2, PGF2α, PGI2) reduced food intake and inhibited AgRP neurons (**Figure 4A–4L**), chemogenetic experiments indicate these effects are variably sensory neuron-dependent (**Figures 5; S5; S6**). Global inhibition of NaV1.8 neurons completely prevented PGE2- and PGD2-induced anorexia **(Figures 5A, 5B)**. By contrast, vagal afferent-specific inhibition partially rescued PGE2 but not PGD2-induced anorexia (**Figures 5I, 5J; S6C, S6G**). PGF2α- and PGI2-induced anorexia were only modestly peripheral sensory neuron-dependent (**Figures 5C, 5D, 5K, 5L**), suggesting these signals may be sensed at the blood-brain barrier or via sensory neuron populations not targeted by our mouse models.

Of note, IL-1β induced anorexia lasts for several hours and dual cyclooxygenase inhibition fully rescues food intake for only around 30 minutes despite ketorolac’s long half-life (**Figure 3A–3C)**^81^. At later time points, ketorolac or selective COX-2 inhibition only partially rescued re-feeding **(Figures 3A–3C; S4G–S4I)**. Collectively, these results suggest that prostaglandin signaling largely in non-vagal afferent neurons is required for the onset of IL-1β-induced anorexia, and that other prostaglandin- and non-prostaglandin-dependent mechanisms sustain anorexia at later time points.

### Prostaglandin-dependent processes mask a role for vagal afferent neurons in mediating prolonged IL-1β-induced anorexia

With respect to prolonged IL-1β-induced anorexia, our studies suggest a role for vagal afferent neurons. Under COX blockade, chemogenetic inhibition of either NaV1.8- or Phox2b-expressing neurons further rescued food intake beyond the effects of ketorolac alone (**Figures 7; S7**), unmasking a prostaglandin-independent component of IL-1β-induced anorexia mediated at least in part by vagal afferent neurons that are likely both NaV1.8- and Phox2b-positive. Our results help reconcile inconsistent findings with respect to vagal involvement in IL-1β-induced anorexia where variations in experimental design, including IL-1β dose, timing of food intake measurements, and route of administration, may determine whether vagal or non-vagal afferent pathways are dominant in driving behavior.

### Limitations of study

Our study has several limitations. Our behavioral studies focus solely on fast re-feeding as a starting point. Future work will be required to understand how IL-1β alters feeding microarchitecture, whether the same circuits regulate *ad libitum* food intake, and whether peripheral sensory neurons are required for the development of IL-1β-induced aversion, as separate CNS circuits mediate anorexia and aversion ^68,82–85^.

Each of the individual mouse models we used in chemogenetic studies has limitations. For example, Phox2b expression is restricted to nodose, petrosal, and cingulate ganglia sensory afferent neurons, but Phox2b-Cre is also expressed in brainstem neurons involved in feeding ^61,62,80^. Similarly, while virus-mediated expression of inhibitory DREADD receptor in NaV1.8:AAV-hM4Di animals is completely vagus-restricted it may result in less robust and less homogeneous neural inhibition than the transgenic models. However, given the generally consistent findings between our two vagus-specific models, including the significant effect of chemogenetic inhibition on CCK-induced anorexia and directionally similar rescue in re-feeding during PGE2-induced anorexia, our results support the involvement of both non-vagal and vagal afferent neurons in IL-1β-induced anorexia.

Finally, we only studied one dose of IL-1β, and some studies suggest that higher concentrations or alternate injection routes of this cytokine may be required to recruit blood-brain barrier signaling ^36,86,87^. More broadly, we only studied pharmacologic effects of inflammatory signals and not the effect of infectious agents or endogenous autoinflammation on behavior and neural circuits.

In addition to these limitations, our findings highlight several other unanswered questions that will be the topic of future studies. The cellular sources and key prostaglandin species driving peripherally induced IL-1β-induced anorexia have yet to be identified ^38^. Finally, the molecular mediators underlying the prostaglandin-independent component of IL-1β-induced anorexia remain unknown. The tools and approaches used here will enable further dissection of these important and complex processes.

## STAR METHODS

### KEY RESOURCES TABLE

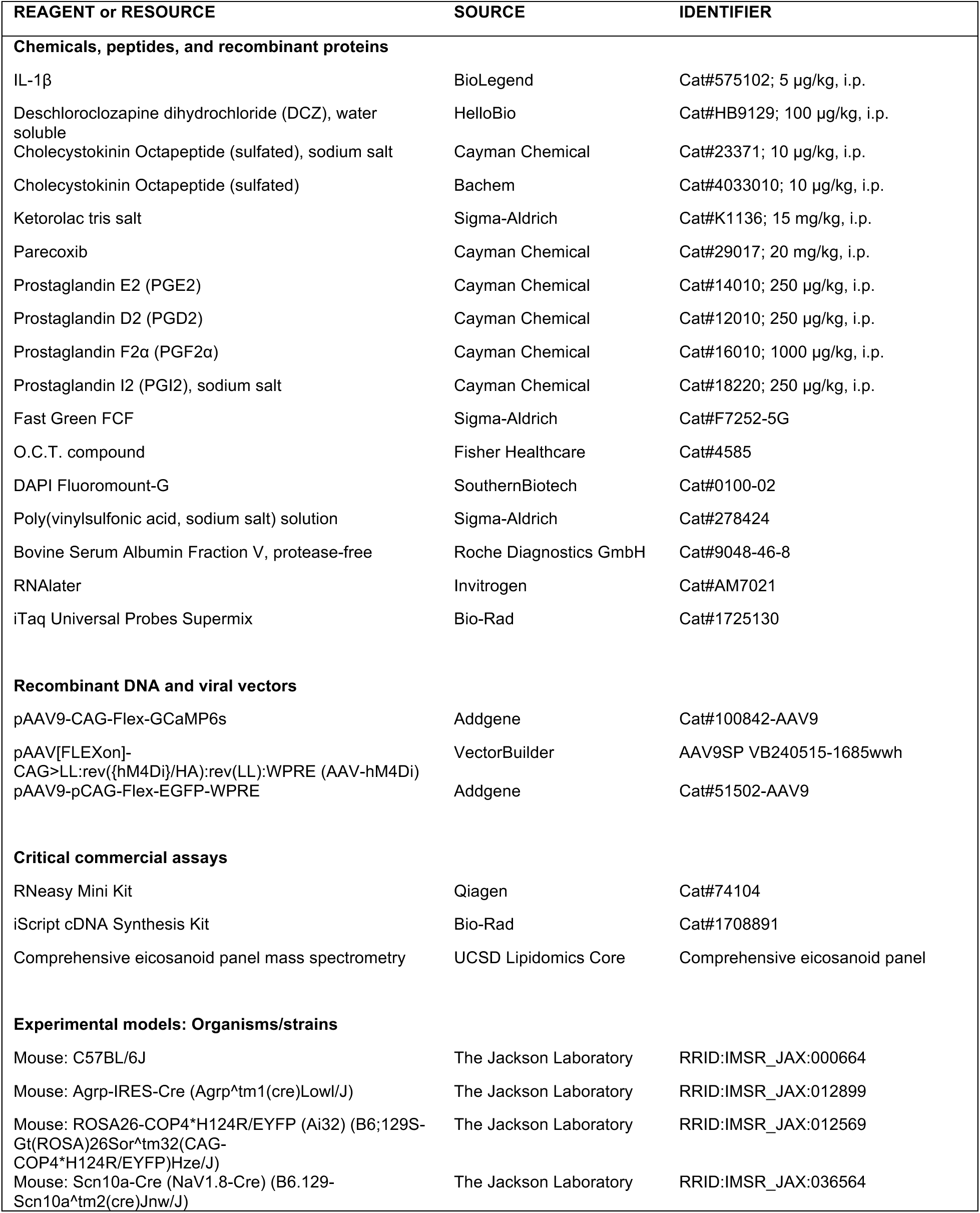
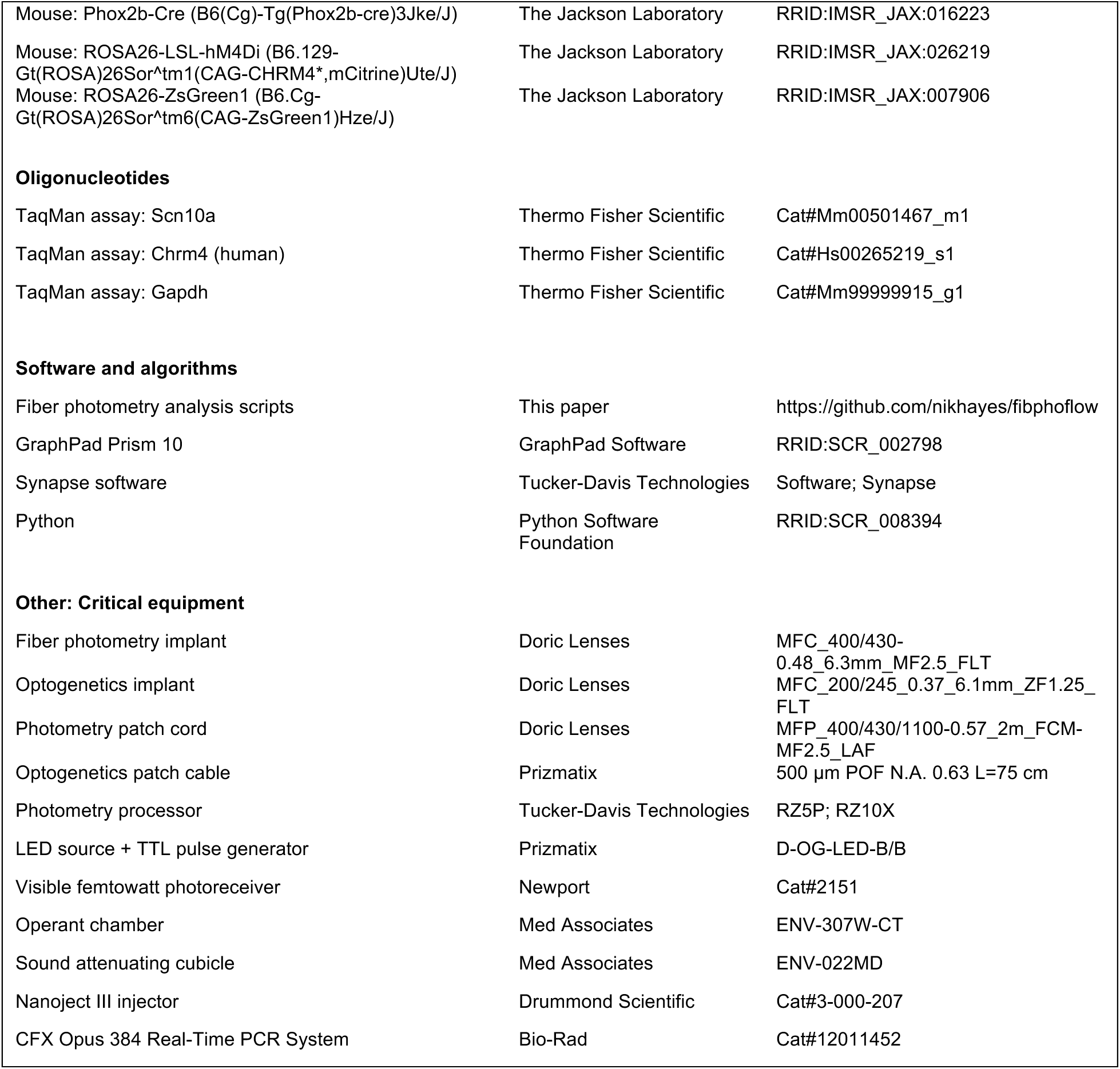

## RESOURCE AVAILABILITY

### Lead contact

Requests for resources and reagents should be directed to the lead contact, Lisa Beutler.

### Materials availability

An AAV vector, pAAV[FLEXon]-CAG>LL:rev({hM4Di}/HA):rev(LL):WPRE (AAV-hM4Di; AAV9SP VB240515-1685wwh, VectorBuilder) was the only new material generated and is available from the investigator or from VectorBuilder.

### Data and code availability

The data reported in this paper will be shared by the lead contact upon request. All original code has been deposited at GitHub (https://github.com/nikhayes/fibphoflow).

## ACKNOWLEDGEMENTS

L.R.B. acknowledges support from the American Diabetes Association Pathway to Stop Diabetes Award (12-22-ACE-31), a McKnight Foundation Neurobiology of Brain Disorders Award, and NIH grants P30-DK020595, K08-DK118188, and R01-DK128477. N.W.H. was additionally supported by NIH grant F31-AI179597.

We thank Dr. Wenfei Han for training in the nodose ganglia injection technique in the de Araujo laboratory.

## AUTHOR CONTRIBUTIONS

N.W.H. and L.R.B. conceptualized the study, designed experiments, and wrote the manuscript. N.W.H. performed experiments and analyzed data. J.L.X., K.K.G., M.J.D., E.T., C.M.L., and J.A.F. performed experiments and assisted with data analysis.

## EXPERIMENTAL MODEL AND SUBJECT DETAILS

Experimental protocols were approved by the Northwestern University IACUC in accordance with NIH guidelines for the Care and Use of Laboratory Animals (Protocol Numbers: IS00021524, IS00015106, IS00023902, IS00011930, IS00016880). Mice were maintained on a 12 h light/12 h dark cycle with *ad libitum* access to standard chow except as specified in the text. All mouse lines were maintained on a C57BL/6J background.

No statistical methods were used to determine sample sizes. The lipidomics experiment (Figure 4) used male mice only while all others used male and female mice aged 10 weeks to 12 months, and data from males and females were combined for analysis. Unless otherwise indicated, experiments involved a 16-hour fast, during which mice had *ad libitum* water access. The experiments were performed during the first half of the dark cycle (Zeitgeber Time ZT12–ZT18) in a reverse light–dark cycle room, and any light exposure was transient and minimized. Mice were grouped-housed unless otherwise specified.

C57BL/6J (000664, Jackson Laboratory) were used to monitor anorexia induced by IL-1β or prostaglandins, as well as for lipidomic experiments. AgRP-Cre mice (Agrp^tm1(cre)Lowl^, 012899, Jackson Laboratory) backcrossed to C57BL/6J were used for fiber photometry studies. For optogenetic studies, AgRP-Cre mice were crossed with ChR2 mice (129S-Gt(ROSA)26Sor^tm^^32(CAG-COP4*H124R/EYFP)Hze^ (ROSA26-LSL-ChR2-eYFP), 012569, Jackson Laboratory) to generate AgRP:ChR2 double-transgenic mice and controls. Chemogenetic manipulation of IL-1β and prostaglandin-induced feeding suppression involved NaV1.8-Cre (B6.129(Cg)-*Scn10a^tm2(cre)Jwo^*/TjpJ, 036564, Jackson Laboratory) and Phox2b-Cre (B6(Cg)-Tg(Phox2b-cre)3Jke/J, 016223, Jackson Laboratory) expressing mice crossed with Cre-dependent hM4Di DREADD mice (B6.129-*Gt(ROSA)26Sor^tm1(CAG-CHRM4*,-mCitrine)Ute^*/J, 026219, Jackson Laboratory) to yield double transgenic NaV1.8:hM4Di or Phox2b:hM4Di mice. All NaV1.8-Cre mice used in experiments were heterozygous at the NaV1.8 locus as homozygous mice are NaV1.8 null. Mice homozygous for the hM4Di allele were used in NaV1.8 experiments whereas Phox2b experiments used mice heterozygous for the hM4Di allele. For experiments where Cre-dependent AAV was injected into the nodose ganglia, NaV1.8-Cre/+ mice with or without an Ai6-ZsGreen allele (B6.Cg-*Gt(ROSA)26Sor^tm6(CAG-ZsGreen1)Hze^*/J, 007906, Jackson Laboratory) were used.

## METHOD DETAILS

### Surgical procedures

**Fiber-photometry cannula implantation:** AgRP-Cre mice were anesthetized with isoflurane and positioned for stereotaxic surgery. Ophthalmic ointment was applied to protect the cornea, and hair over the skull was removed using depilatory cream. Body temperature was maintained via a heating pad. Post-operatively, mice received sustained-release buprenorphine (1.5 mg/kg) and meloxicam (2 mg/kg) for analgesia.

A craniotomy was made at x = +0.26 mm, y = −1.35 mm relative to bregma using a dental drill. A pulled glass pipette delivered 400 nL pAAV-CAG-Flex-GCaMP6s (100842-AAV9, Addgene) at z = −5.95 mm (50 nL/min; Nanoject III injector, Drummond Scientific). A fiber cannula (MFC_400/430-0.48_6.3mm_MF2.5_FLT, Doric Lenses) was implanted at the same coordinates and secured with dental cement. A bronze mating sleeve (SLEEVE_BR_2.5, Doric Lenses) was also cemented in place. Mice were allowed to recover at least two weeks to allow for viral expression before initial photometry experiments.

**Optogenetics cannula implantation:** AgRP-Cre/+;ROSA26-LSL-ChR2-eYFP (AgRP:ChR2) mice and Cre-negative ROSA26-LSL-ChR2-eYFP (ChR2 Ctrl) littermate controls were prepared for stereotaxic injection and fiber implantation as above, and also received the same post-operative care. For the optogenetic cannula implantation, an optical fiber (MFC_200/245_0.37_6.1mm_ZF1.25_FLT, Doric Lenses) was unilaterally implanted above the arcuate nucleus (x = +0.26 mm, y = −1.35 mm, z = −5.80 mm from bregma) and secured with dental cement. Mice were allowed to recover for at least one week before optogenetic experiments.

**Bilateral nodose ganglia injections:** Adapted from Han et al.^66^ with minor modifications to equipment and materials. Briefly, aluminosilicate glass pipettes (A100-64-10, Sutter) were pulled on a P-97 puller, tip-trimmed, and beveled at a 30° angle to achieve a ∼15 µm outer diameter (EG-402, Narishige). Pipettes were backfilled with viral solution containing 0.02 mg/mL Fast Green FCF (F7252-5G, Sigma) to visualize injection sites.

Mice were anesthetized with isoflurane, ophthalmic ointment was applied, and the neck was depilated. A ∼1.5 cm midline incision was made to expose each nodose ganglion sequentially. Using a Nanoject III injector (3-000-207, Drummond Scientific), 600 nL of AAV-hM4Di (AAV9SP VB240515-1685wwh, VectorBuilder) or AAV-Ctrl (51502-AAV9, Addgene) was delivered into each ganglion at 100 nL/min. The incision was closed with 6-0 polypropylene sutures (S-P618R13, AD Surgical).

Postoperatively, mice received meloxicam and buprenorphine as above and were allowed to recover for at least 3 weeks before experimentation.

### Fiber-photometry recordings

Two separate fiber photometry configurations were used for data acquisition. In the first configuration, the photometry processor (RZ5P, Tucker-Davis Technologies) was separate from the LED driver (DC4100, Thorlabs) and LEDs (M405FP1 and M470F3, Thorlabs); in the second configuration, an integrated unit (RZ10X, Tucker-Davis Technologies) housed the processor, LED driver, and LEDs. For each experimental mouse, the same photometry system and patch cord (MFP_400/430/1100-0.57_2m_FCM-MF2.5_LAF, Doric Lenses) were used across all recording sessions to allow reliable within-mouse comparisons of calcium signals over time.

Mice were placed in operant chambers (ENV-307W-CT, Med Associates) housed inside sound-attenuating cubicles (ENV-022MD, Med Associates) in a dedicated dark room isolated from other experiments. Recordings were conducted with no food or water access unless stated otherwise.

Continuous blue (465–470 nm, GCaMP channel) and UV (405 nm, isosbestic channel) LEDs were each sinusoidally modulated at distinct carrier frequencies. Light was passed through a filtered minicube (Doric Lenses) and delivered via 400/430 µm patch cords to fiber photometry implants. Emitted GCaMP6s and isosbestic signals traveled back along the same path to a femtowatt photoreceiver (model 2151, Newport) or the integrated Lux photosensors (RZ10X, Tucker-Davis Technologies). Signals were sampled at 1.0173 kHz, demodulated by lock-in amplification, and recorded by the TDT processor (RZ5P or RZ10X) using Synapse software (Tucker-Davis Technologies). The raw data were processed with custom Python scripts (https://github.com/nikhayes/fibphoflow).

Subjects failing to exhibit a baseline ΔF/F AgRP neuron calcium response of ≥20% to chow presentation were considered technical failures excluded from experiments ^52,88^. One to two trials per mouse were averaged as a single biological replicate.

### Feeding experiments

Mice underwent at least five 1-hour habituation sessions to handling, i.p. phosphate buffered saline (PBS) vehicle injections, and temporary isolation from cage mates while in feeding chambers. At least two of these sessions involved overnight fasting then re-feeding during the habituation session to simulate experimental conditions.

**Standard fast re-feeding protocol:** Mice were fasted 16 hours with at least 3 days between experiments involving PBS (vehicle) or cholecystokinin octapeptide (CCK) i.p. injections, and at least 7 days between experiments involving any inflammatory substances. Re-feeding experiments were conducted within the first 6 hours of the dark phase in reverse light–cycle rooms. Mice were placed in their experimental cage and given at least 30 minutes of re-acclimation before re-feeding, which included the time required for one or two sequential i.p. injections. In experiments with two injections, mice received the first injection followed 20 minutes later by a second injection, with exact combinations and dosages indicated in the figures. The following substances were administered: PBS, DCZ (100 µg/kg), ketorolac (15 mg/kg), IL-1β (5 µg/kg), CCK (10 µg/kg), PGE2 (250 µg/kg), PGD2 (250 µg/kg), PGF2α (1 mg/kg), or PGI2 (250 µg/kg).

For our initial IL-1β–specific experiments (Figure 1), re-feeding was performed immediately after injection, whereas later experiments involving IL-1β included a 15-minute interval between the IL-1β injection and re-feeding to allow IL-1β’s anorectic effects to manifest. For experiments involving prostaglandins or CCK, mice were re-fed immediately after injection. Two standard chow pellets were placed in each mouse’s wire rack, and the weight was measured every 15 minutes for the duration indicated in the experimental figures. Mice eating < 0.3 g in the first hour of control PBS-injected re-feeding experiments were excluded from studies.

**Optogenetic feeding protocol**: Optogenetic experiments were performed in AgRP:ChR2 mice and ChR2 control mice. Mice were habituated as above, and to tethering to optogenetic patch cords (500µm POF N.A. 0.63 L=75cm, Prizmatix) across at least five 1-hour sessions, including a minimum of two sessions that involved overnight fasting and re-feeding to simulate experimental conditions.

Blue light stimulation (460 nm, 2 seconds ON/3 seconds OFF, 10 ms pulse width, 20 Hz, 10–20 mW) was generated using an LED source and programmable TTL pulse generator (D-OG-LED-B/B, Prizmatix). Fiber-optic patch cords were connected to the implanted optical fibers via zirconia mating sleeves (F210-3001, Doric Lenses).

On experimental days, mice were fasted overnight (16 hours) with water access, then placed individually into behavior chambers without food or water. Each session began with 30 minutes of habituation without light stimulation. Mice then received an intraperitoneal injection of either PBS or IL-1β (5 µg/kg; 575102, BioLegend) and remained in the chamber for 15 minutes before re-feeding. At the start of re-feeding, a single pre-weighed chow pellet was placed in the food rack, and 30 minutes of food intake was recorded.

Optogenetic stimulation was initiated simultaneously with food presentation in stimulated sessions. Stimulated and non-stimulated sessions were conducted in the same mice on separate days in a counterbalanced order, with at least 3 days between sessions for PBS treatment and a minimum of 7 days between any sessions involving IL-1β. Only AgRP:ChR2 mice that exhibited at least a twofold increase in fast re-feeding during optogenetic stimulation following PBS injection compared to their non-stimulated PBS control session were included in the analyses as previously described ^52,88^.

**DREADD chemogenetic experiments:** Mice included NaV1.8-Cre;hM4Di/hM4Di (NaV1.8:hM4Di), NaV1.8-Cre heterozygous controls (NaV1.8:Ctrl), Phox2b-Cre;hM4Di/+ (Phox2b:hM4Di), Phox2b-Cre controls (Phox2b:Ctrl), and NaV1.8-Cre heterozygous mice injected bilaterally with AAV9-CAG-Flex-hM4Di-HA (NaV1.8:AAV-hM4Di) or AAV9-CAG-Flex-EGFP (NaV1.8:AAV-Ctrl) in the nodose ganglia. On experimental days, mice received i.p. injections of vehicle (PBS), deschloroclozapine dihydrochloride (DCZ), or DCZ+Ketorolac 20 minutes before a second i.p. injection of CCK (10 µg/kg), IL-1β (5 µg/kg), prostaglandins (PGE2, 250 µg/kg; PGD2, 250 µg/kg; PGF2α, 1 mg/kg; or PGI2, 250 µg/kg), or PBS vehicle. Treatments were counter-balanced such that half of each experimental cohort received PBS vehicle while the other half received DCZ on a given day, with the assignments reversed on the subsequent experimental day.

### Plasma collection and lipidomic measurements

C57BL/6J mice were habituated as described for feeding experiments and fasted for 16 hours with *ad libitum* water access. On the day of plasma collection, mice received two sequential intraperitoneal injections 20 min apart: PBS followed by PBS, PBS followed by IL-1β (5 µg/kg), or ketorolac (15 mg/kg) followed by IL-1β (5 µg/kg). Mice were briefly anesthetized with isoflurane 30 minutes after the second injection then euthanized by cervical dislocation. Blood was collected by cardiac puncture into EDTA-coated tubes on ice. Plasma was separated at 4°C by centrifugation (1,500×g for 15 min) and 50 µL aliquots were snap-frozen in liquid nitrogen to minimize ex vivo eicosanoid production. Samples were stored and shipped at −80°C to the UCSD Lipidomics Core for comprehensive eicosanoid panel mass spectrometry.

### Histology & imaging

NaV1.8-Cre/+ mice that had been injected with AAV-Ctrl, which encodes Cre-dependent EGFP, were perfused transcardially with PBS then 4% paraformaldehyde. Nodose-Jugular-Petrosal ganglia were post-fixed for 24 h, washed with PBS, cryoprotected in 30% sucrose overnight, and flash frozen in O.C.T. compound (Fisher Healthcare). Nodose ganglia were cryosectioned at 20 µm (CM1860, Leica). Sections were mounted with Fluoromount-G (0100-02, SouthernBiotech) and EGFP fluorescence in transduced NaV1.8-positive nodose neurons was imaged on a Leica Thunder Tissue Imager.

### RNA isolation

Nodose ganglia were harvested from NaV1.8-Cre/+ mice that had been injected into the nodose ganglia bilaterally with AAV-hM4Di or AAV-Ctrl and had completed feeding experiments.

Dissection was performed as described by Han and de Araujo ^66^. Mice were briefly anesthetized with isoflurane and then euthanized by decapitation. Nodose ganglia were dissected in an ice-cold solution of phosphate-buffered saline (PBS, Ca²⁺/Mg²⁺-free) containing 0.5% poly(vinylsulfonic acid) (278424, Sigma Aldrich) and 0.1% bovine serum albumin (9048-46-8, Roche Diagnostics) to minimize RNase activity and prevent tissue adhesion. Bilateral ganglia dissections were completed within fifteen minutes post euthanasia and then placed in room temperature RNAlater (AM7021, Invitrogen) for at least 30 minutes before storing overnight at 4°C.

Nodose ganglia were homogenized with a glass pestle homogenizer in 350 µL Buffer RLT Plus (Qiagen), and RNA was isolated using the RNeasy Mini Kit (74104, Qiagen) according to the manufacturer’s instructions.

### qRT-PCR

RNA was reverse transcribed using the iScript cDNA Synthesis Kit (1708891, Bio-Rad) following the manufacturer’s instructions. The resulting cDNA was diluted and used for quantitative PCR with the iTaq Universal Probes Supermix (1725130, Bio-Rad) on a CFX Opus 384 Real-Time PCR System (12011452, Bio-Rad). TaqMan Gene Expression Assays with FAM-MGB probes were used for target detection. Each sample was run in triplicate for every probe mix.

Expression of *Scn10a* and *CHRM4* (the human hM4Di receptor component) was quantified using the relative standard curve method and normalized to the housekeeping gene *Gapdh*.

Thermo Fisher Scientific TaqMan assay probe IDs for each target are listed in Key Resources Table.

### Quantification and Statistical Analysis

All statistical analyses were performed using GraphPad Prism 10. Sample sizes are specified in figure legends, and all tests were two-tailed unless otherwise stated. Adjusted p-values were reported where multiple comparisons were performed using the Holm–Šídák correction method.

**Photometry analysis:** Custom Python scripts (https://github.com/nikhayes/fibphoflow) were used to preprocess, analyze, and visualize fiber-photometry recordings. AgRP neuron GCaMP fluorescence (465–470 nm excitation) and the corresponding isosbestic control (405 nm excitation) were despiked using a 100-ms non-overlapping block-median filter, then anti-aliased and downsampled to 1 Hz. The resulting traces were smoothed using a low-pass filter; all subsequent normalization and analyses were performed on these smoothed traces.

For stimulus-evoked responses, each channel was expressed as percent ΔF/F relative to a pre-injection baseline (F₀), where F₀ was defined as the mean fluorescence during a 5-min baseline window preceding injection (8 to 3 min prior to stimulus): ΔF/F = (Fₜ/F₀ − 1) × 100. To reduce shared artifacts (e.g., motion and slow fluorescence fluctuations), a corrected calcium-dependent signal was computed by subtracting the baseline-normalized isosbestic (405 nm) ΔF/F trace from the baseline-normalized GCaMP (470 nm) ΔF/F trace for data visualization.

AUC values derived from photometry traces were compared between treatments using two-way ANOVA when multiple time windows and treatment factors were present or paired two-tailed t-tests when comparing a single condition within subjects.

**Food intake analysis:** Standard chow was manually weighed at the specified time points during feeding experiments. Cumulative intake was calculated by summing the reductions in pellet weight across successive time points.

Feeding data were analyzed by two-way repeated-measures ANOVA. When appropriate, the Geisser–Greenhouse correction was applied for violations of sphericity, and significant main effects or interactions were followed by Holm–Šídák post hoc tests.

**Lipidomic analysis:** Plasma eicosanoid concentrations were measured by comprehensive eicosanoid panel mass spectrometry (UCSD Lipidomics Core). Values reported as non-detectable were treated as below the assay limit of detection (LOD). For analyses requiring numeric values, non-detectable values were imputed as one-fifth of the minimum detected concentration for that analyte across all samples, following UCSD Lipidomics Core guidance. Detection frequencies were compared using one-tailed Fisher’s exact tests on detection status given the expected increase in prostaglandins after IL-1β administration, with Holm–Šidák correction applied for multiple planned comparisons.

**qPCR analysis:** RNA extracted from nodose ganglia was reverse-transcribed and quantified by RT-qPCR using TaqMan assays, with expression calculated as 2^–ΔCt relative to *Gapdh*.

Samples without detectable amplification by 40 cycles were assigned Ct = 40 prior to ΔCt calculation; “detected” was defined as Ct < 40. For *Scn10a* (NaV1.8), all samples were detected, and relative expression between groups was compared using the Mann–Whitney test. For *CHRM4* (hM4Di transgene), the NaV1.8:AAV-Ctrl group exhibited complete non-detection, so detection frequencies were compared using Fisher’s exact test. Open symbols in figures denote non-detected samples.

---

**Figure S1.**
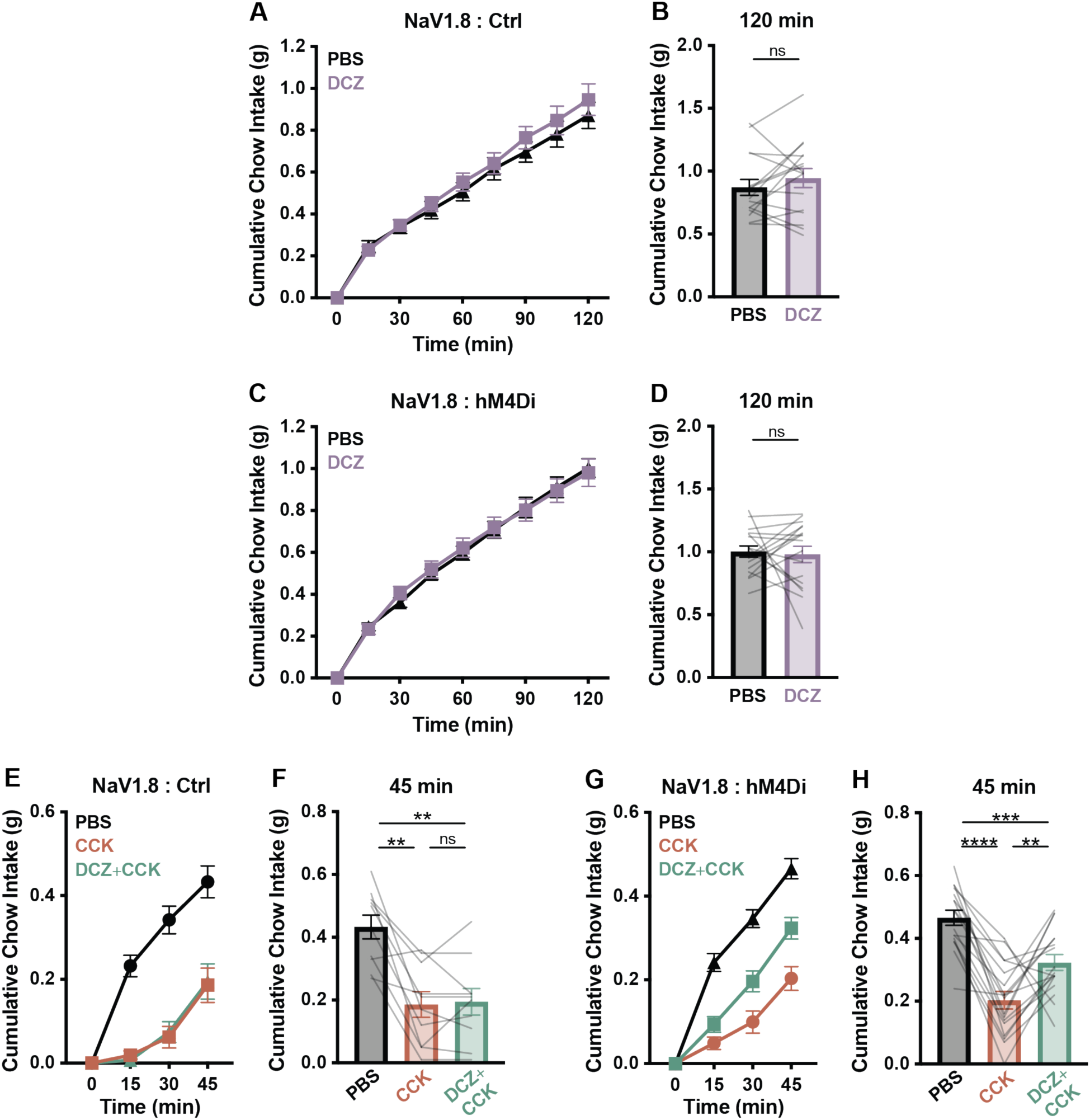
NaV1.8-expressing peripheral sensory neuron inhibition does not impact fast re-feeding and partially rescues CCK-induced anorexia. **(A–D)** Cumulative chow intake at the indicated time points in fasted NaV1.8:Ctrl **(A, B)** and NaV1.8:hM4Di **(C, D)** mice following i.p. injection with PBS or DCZ (100 μg/kg). **(A)** Two-way repeated-measures ANOVA: main effect of time, p < 0.0001; main effect of treatment, p = 0.33; time×treatment interaction p = 0.22. n = 16, paired design. **(C)** Two-way repeated-measures ANOVA: main effect of time, p < 0.0001; main effect of treatment, p = 0.92; time×treatment interaction, p = 0.43. n = 17 mice, paired design. **(E–H)** Cumulative chow intake at the indicated time points in fasted NaV1.8:Ctrl (E, F) and NaV1.8:hM4Di **(G, H)** mice following i.p. injection with PBS, CCK (10 μg/kg), or DCZ (100 μg/kg)+CCK. **(E)** Two-way repeated-measures ANOVA: main effects of time, treatment, and time×treatment, p < 0.0001, n = 10 mice, paired design. **(G)** Two-way repeated-measures ANOVA: main effects of time, treatment, and time×treatment, p < 0.0001. n = 17 mice, paired design. **(A–H)** Error bars indicate mean ± SEM. Lines represent individual mice. Holm–Šídák post hoc comparisons: **p<0.01, ***p<0.001, ****p<0.0001

**Figure S2.**
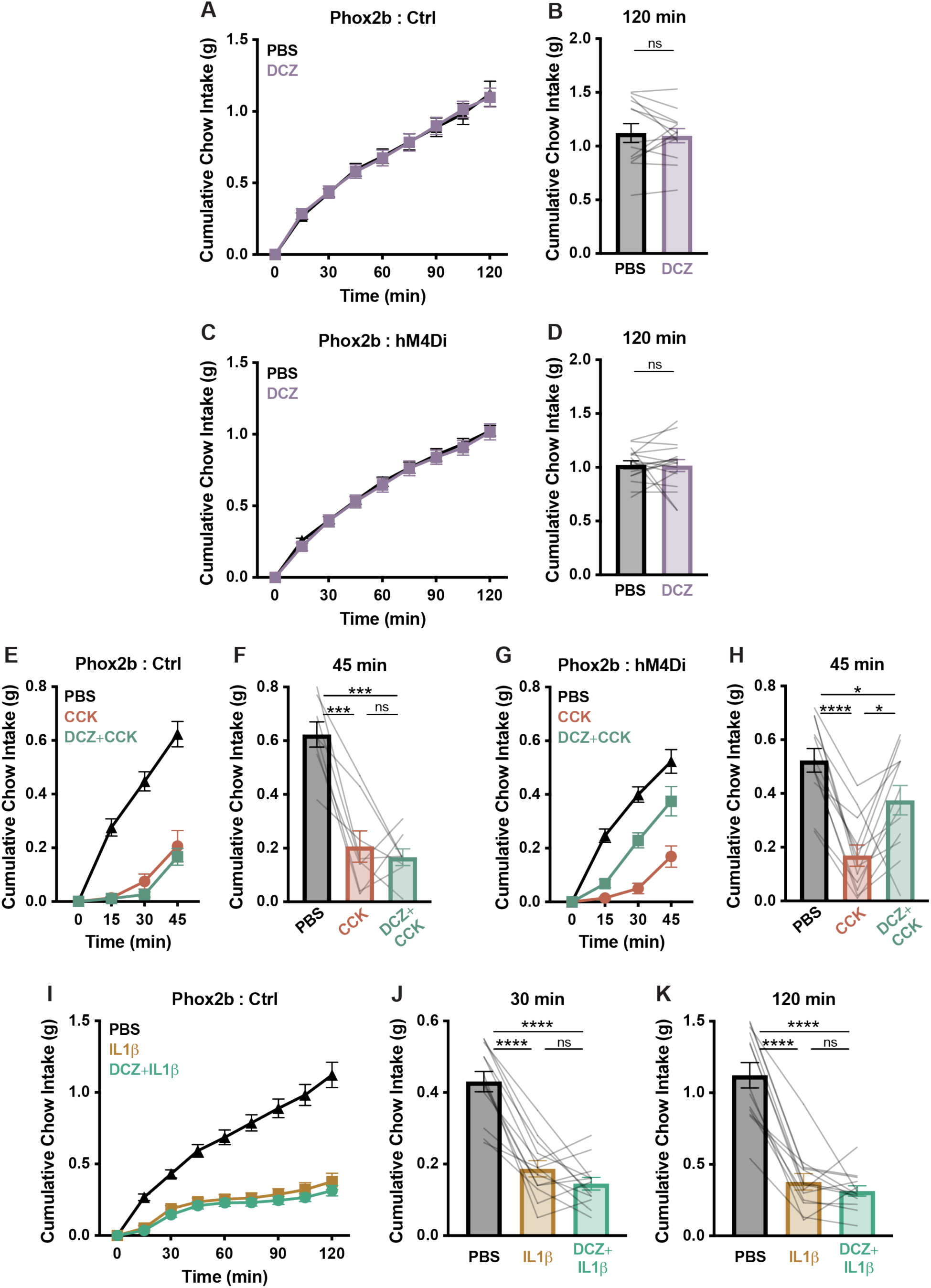
Inhibition of Phox2b-expressing vagal afferent neurons does not impact fast re-feeding and partially rescues CCK-induced anorexia. **(A–D)** Cumulative chow intake at the indicated time points in fasted Phox2b:Ctrl (A, B) and Phox2b:hM4Di **(C, D)** mice following i.p. injection with PBS or DCZ (100 μg/kg). **(A)** Two-way repeated-measures ANOVA: main effect of time, p < 0.0001; main effect of treatment, p = 0.96; time×treatment interaction p = 0.67. n = 13 mice, paired design. **(C)** Two-way repeated measures ANOVA: main effect of time, p < 0.0001; main effect of treatment, p = 0.75; time×treatment interaction, p = 0.84. n = 17 mice, paired design. **(E–H)** Cumulative chow intake at the indicated time points in fasted Phox2b:Ctrl (E, F) and Phox2b:hM4Di **(G, H)** mice following i.p. injection with PBS, CCK (10 μg/kg), or DCZ (100 μg/kg)+CCK. **(E)** Two-way repeated-measures ANOVA: main effects of time, treatment, and time×treatment, p < 0.0001. n = 8 mice, paired design. **(G)** Two-way repeated-measures ANOVA: main effects of time, treatment, and time×treatment, p < 0.0001. n = 12 mice, paired design. **(I–K)** Cumulative chow intake at the indicated time points in fasted Phox2b:Ctrl mice following i.p. injection of PBS, IL-1β (5 μg/kg), or DCZ (100 μg/kg)+IL-1β. **(I)** Two-way repeated-measures ANOVA: main effects of time, treatment, and time×treatment interaction, p < 0.0001. n = 13 mice, paired design. **(A–K)** Error bars indicate mean ± SEM. Lines represent individual mice. Holm–Šídák post hoc comparisons: **p<0.01, ***p<0.001, ****p<0.0001

**Figure S3.**
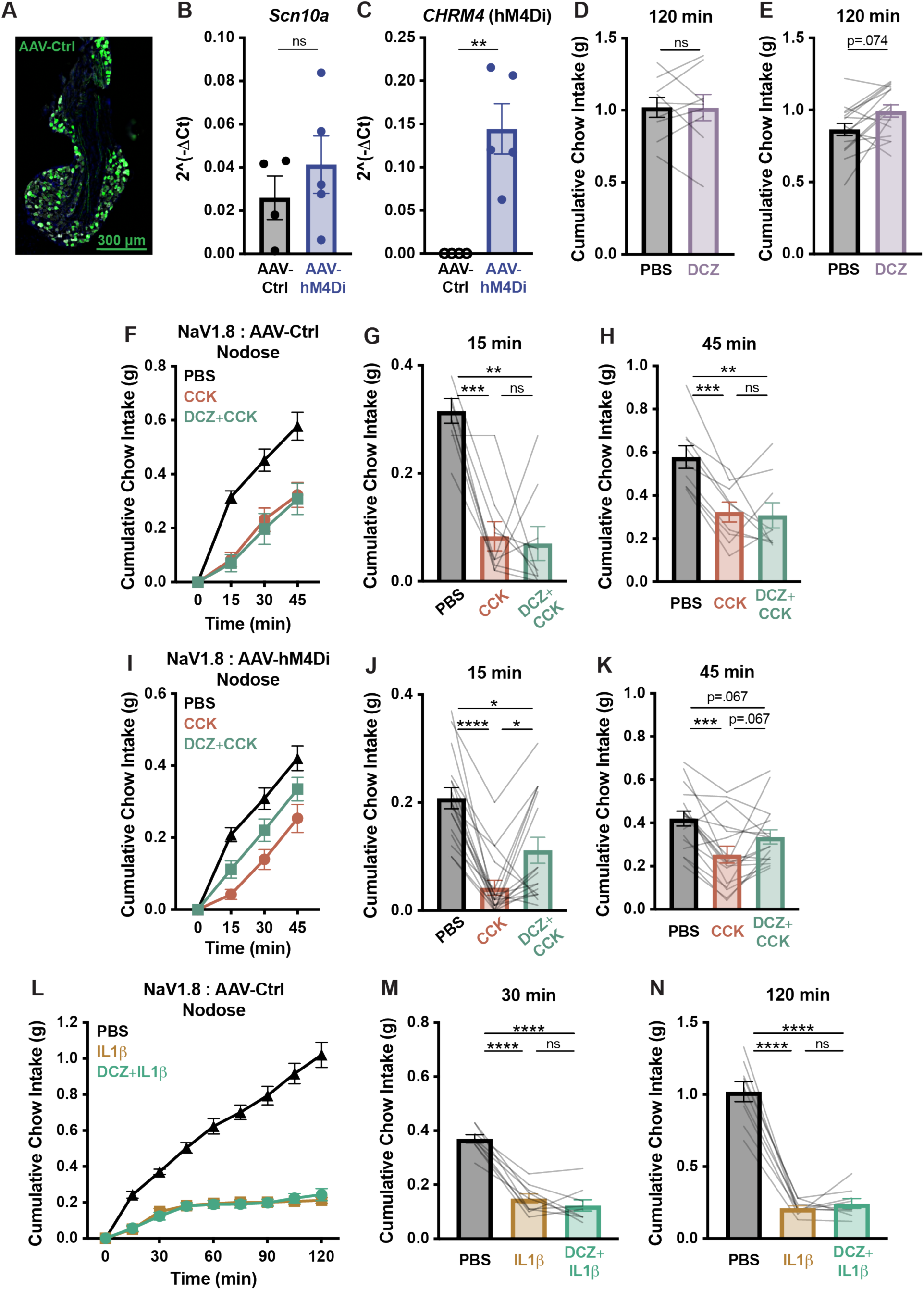
Inhibiting NaV1.8-expressing nodose ganglia neurons via virus-mediated DREADD expression enhances fast re-feeding and partially rescues CCK-induced anorexia. **(A)** AAV-Ctrl-mediated EGFP expression in the nodose ganglion of a NaV1.8-Cre/+ (NaV1.8:AAV-Ctrl) mouse. **(B, C)** Scn10a **(B)** and CHRM4 **(C)** mRNA expression in nodose ganglia measured by RT-qPCR (2^–ΔCt values normalized to Gapdh) in NaV1.8-Cre mice injected bilaterally into nodose ganglia with AAV-Ctrl (n = 4) or AAV-hM4Di (n = 5). **(B)** Mann–Whitney test, p = 0.557. **(C)** Two-tailed Fisher’s exact test, adjusted p = 0.0079. Filled symbols indicate detectable values, hollow symbols indicate non-detected amplification (set at limit of detection). **(D, E)** Cumulative chow intake at the indicated time point in fasted NaV1.8:AAV-Ctrl **(D)** or NaV1.8:AAV-hM4Di **(E)** nodose mice after i.p. injection of PBS or DCZ (100 μg/kg). **(D)** Two-way repeated-measures ANOVA: main effect of time, p < 0.0001; main effect of treatment, p = 0.95; time×treatment interaction, p = 0.76. n = 9 mice, paired design. **(E)** Two-way repeated-measures ANOVA: main effect of time, p < 0.0001; main effect of treatment, p = 0.0187; time×treatment interaction, p = 0.0095. n = 16 mice, paired design. **(F–K)** Cumulative chow intake at the indicated time points in fasted NaV1.8:AAV-Ctrl **(F–H)** or NaV1.8:AAV-hM4Di nodose mice **(I–K)** following i.p. injection of PBS, CCK (10 μg/kg), or DCZ (100 μg/kg)+CCK. **(F)** Two-way repeated-measures ANOVA: main effect of time, p < 0.0001; main effect of treatment, p = 0.0014; time×treatment interaction, p = 0.0003. n = 9 mice, paired design. **(I)** Two-way repeated measures ANOVA: main effects of time, treatment, and time×treatment interaction, p < 0.0001. n = 17 mice, paired design. **(L–N)** Cumulative chow intake at the indicated time points in fasted NaV1.8:AAV-Ctrl mice following i.p. injection of PBS, IL-1β (5 μg/kg), or DCZ (100 μg/kg)+IL-1β. **(L)** Two-way repeated measures ANOVA: main effects of time, treatment, and time×treatment interaction, p < 0.0001. n = 9 mice, paired design. **(B–N)** Error bars indicate mean ± SEM. Lines represent individual mice. Holm–Šídák post hoc comparisons: *p<0.05, **p<0.01, ***p<0.001, ****p<0.0001

**Figure S4.**
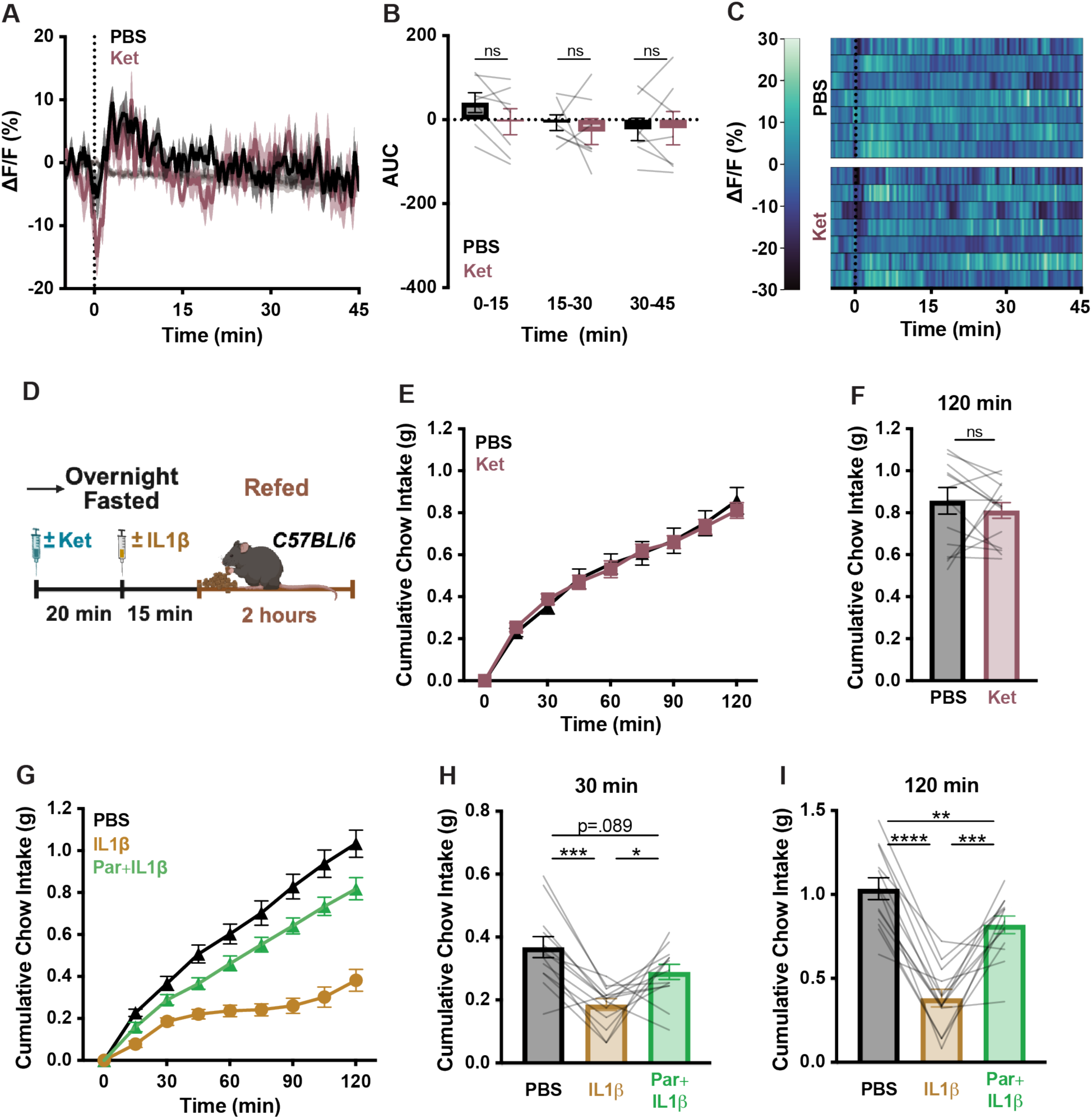
COX inhibition alone does not alter AgRP neuron activity or food intake. (A) AgRP neuron calcium signal in fasted mice injected i.p. with PBS or ketorolac (Ket, 15 mg/kg) at time 0. Isosbestic traces are shown in gray, vertical dashed lines indicate injection time, and traces represent mean ± SEM. n = 7 mice, paired design. (B) Area under the curve (AUC) of traces from **(A)** across 3 15-min time windows following injection. Two-way repeated-measures ANOVA: main effect of time window, p = 0.2045; main effect of treatment, p = 0.4337; time×treatment interaction, p = 0.1292. (C) Heatmaps of ΔF/F (%) in individual mice from **(A)**, with subject order preserved across conditions. (D) Injection and fast re-feeding timing for Figures 3A–3C and S4E, S4F. **(E, F)** Cumulative chow intake at the indicated time points in fasted C57BL/6 mice following i.p. injection with PBS or ketorolac (15 mg/kg). Two-way repeated-measures ANOVA: main effect of time, p < 0.0001; main effect of treatment, p = 0.9164; time×treatment interaction, p = 0.2421. n = 15 mice, paired design. **(G–I)** Cumulative chow intake at the indicated time points in fasted C57BL/6 mice following i.p. injection with PBS, IL-1β (5 μg/kg), or parecoxib (Par, 20 mg/kg)+IL-1β. Two-way repeated-measures ANOVA: main effects of time, treatment, and time×treatment interaction, p < 0.0001. n = 13 mice, paired design. **(B, E, F, G, H, I)** Error bars indicate mean ± SEM. Lines represent individual mice. Holm–Šídák post hoc comparisons: *p<0.05, **p<0.01, ***p<0.001, ****p<0.0001

**Figure S5.**
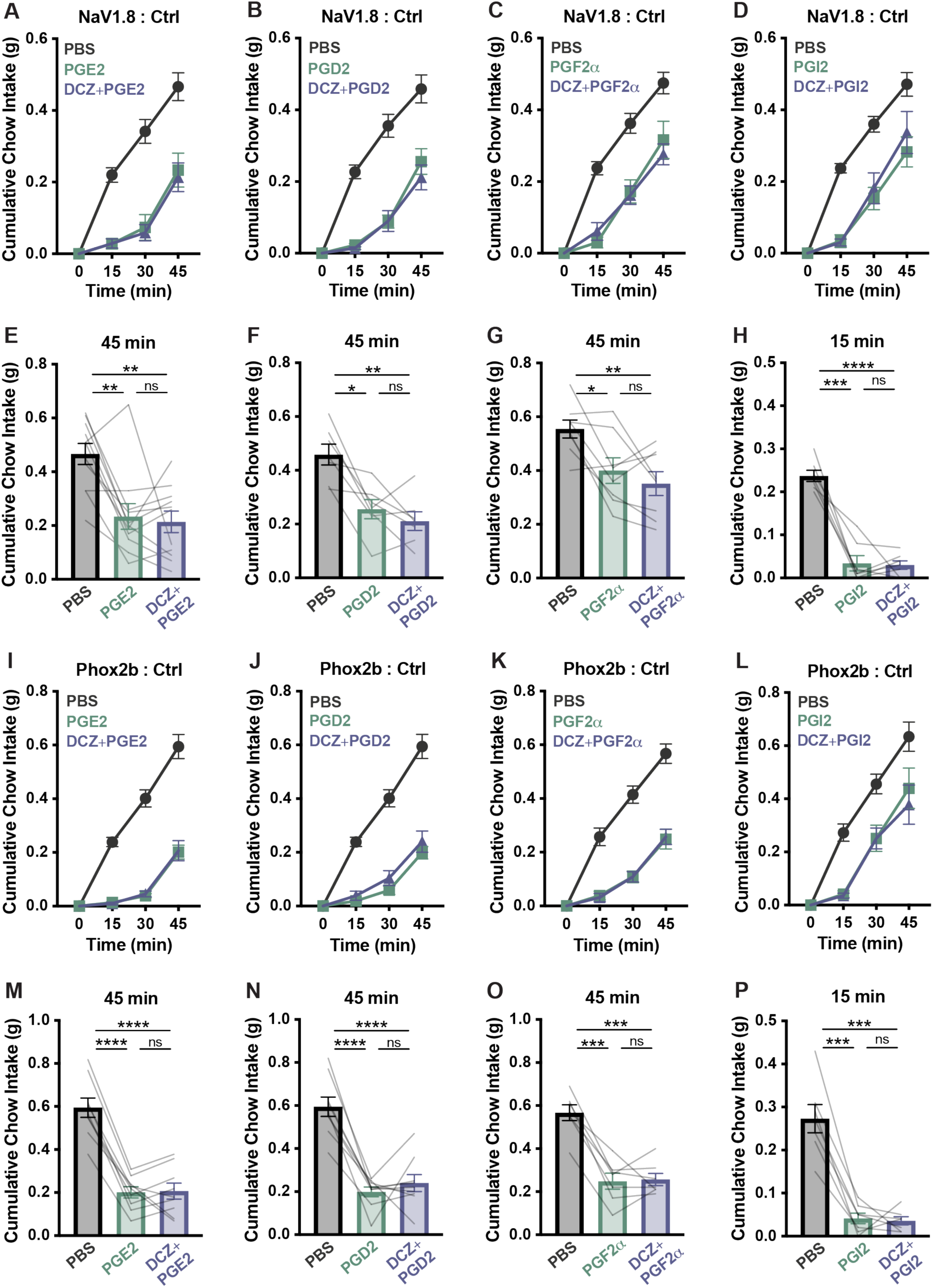
DCZ has no effect on prostaglandin-induced anorexia in NaV1.8-Cre or Phox2b-Cre-expressing control animals. **(A–H)** Cumulative chow intake at the indicated time points in fasted NaV1.8:Ctrl mice following i.p. injection with PBS, PGE2 (250 μg/kg) **(A, E)**, PGD2 (250 μg/kg) **(B, F)**, PGF2α (1 mg/kg) **(C, G)**, or PGI2 (250 μg/kg) **(D, H)**, with or without DCZ (100 μg/kg) pretreatment. **(A–C)** Two-way repeated-measures ANOVA: main effects of time, treatment, and time×treatment interaction, p < 0.0001. **(D)** Two-way repeated-measures ANOVA: main effect of time, p < 0.0001; main effect of treatment, p = 0.0012; time×treatment interaction, p = 0.0025. n = 7-11 mice, paired design. **(I–P)** Cumulative chow intake at the indicated time points in fasted Phox2b:Ctrl mice following i.p. injection with PBS, PGE2 (250 μg/kg) **(I, M)**, PGD2 (250 μg/kg) **(J, N)**, PGF2α (1 mg/kg) **(K, O)**, or PGI2 (250 μg/kg) **(L, P)**, with or without DCZ (100 μg/kg) pretreatment. **(I, J, K)** Two-way repeated-measures ANOVA: main effects of time, treatment, and time×treatment interaction, p < 0.0001. **(L)** Two-way repeated-measures ANOVA: main effect of time, p = 0.0002; main effect of treatment, p = 0.0003; time×treatment interaction, p = 0.0008. n = 7-9 mice, paired design. **(A–P)** Error bars indicate mean ± SEM. Lines represent individual mice. Holm–Šídák post hoc comparisons: *p<0.05, **p<0.01, ***p<0.001, ****p<0.0001.

**Figure S6.**
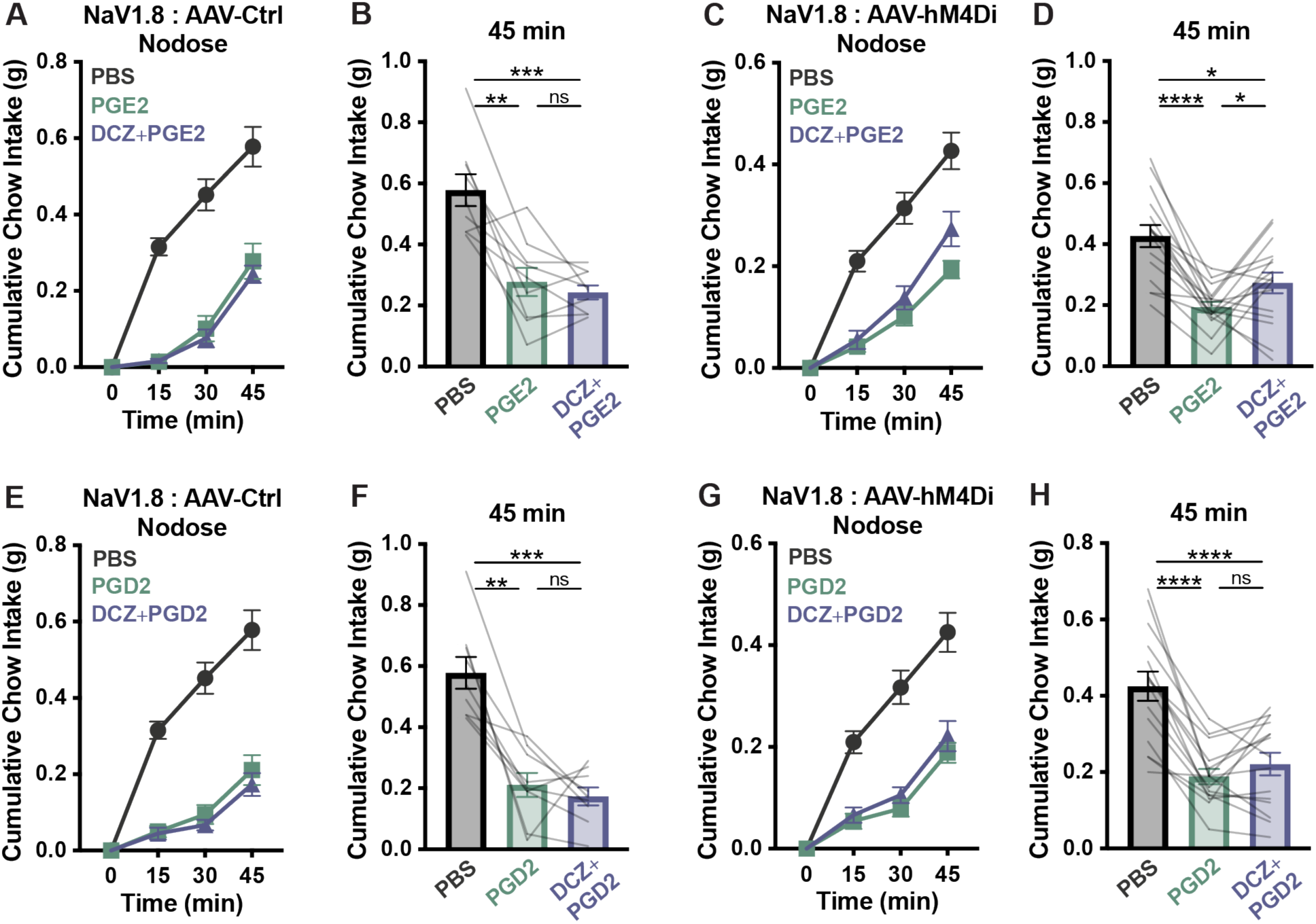
Inhibiting NaV1.8-expressing nodose ganglia neurons via virus-mediated DREADD expression has a limited effect on prostaglandin-induced anorexia. **(A–H)** Cumulative chow intake at the indicated time points in fasted NaV1.8:AAV-Ctrl **(A, B, E, F)** or NaV1.8:AAV-hM4Di **(C, D, G, H)** nodose mice following i.p. injection with PBS, PGE2 (250 μg/kg**) (A–D)** or PGD2 (250 μg/kg) **(E–H)**, with or without DCZ (100 μg/kg) pretreatment **(A, C, E, G)**. Two-way repeated-measures ANOVA: main effects of time, treatment, and time×treatment interaction, p < 0.0001. n = 9–16 mice, paired design. **(A–H)** Error bars indicate mean ± SEM. Lines represent individual mice. Holm–Šídák post hoc comparisons: *p<0.05, **p<0.01, ***p<0.001, ****p<0.0001

**Figure S7.**
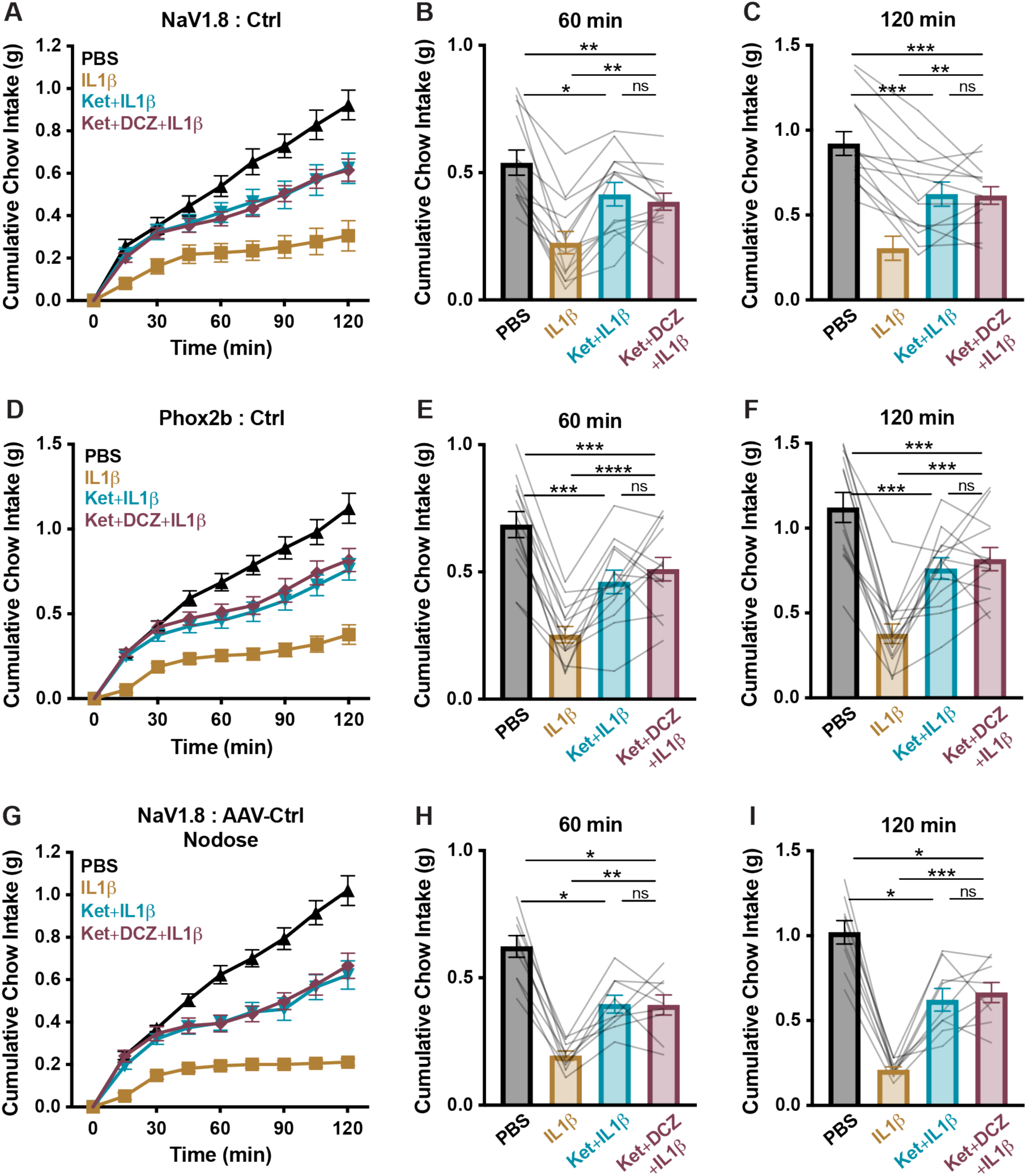
DCZ does not restore the prostaglandin-independent component of IL-1β-induced anorexia in control mice. **(A–C)** Cumulative chow intake at the indicated time points in fasted NaV1.8:Ctrl mice following i.p. injection with PBS, IL-1β (5 μg/kg), ketorolac (Ket, 15 mg/kg)+IL-1β, or DCZ (100 μg/kg)+ketorolac+IL-1β. **(A)** Two-way repeated-measures ANOVA: main effects of time, treatment, and time×treatment interaction, p < 0.0001. n = 13 mice, paired design. **(D–F)** Cumulative chow intake at the indicated time points in fasted Phox2b:Ctrl mice following i.p. injection with PBS, IL-1β (5 μg/kg), ketorolac (15 mg/kg)+IL-1β, or DCZ (100 μg/kg)+ketorolac+IL-1β. **(D)** Two-way repeated-measures ANOVA: main effects of time, treatment, and time×treatment interaction, p < 0.0001. n = 13 mice, paired design. **(G–I)** Cumulative chow intake at the indicated time points in fasted NaV1.8:AAV-Ctrl nodose mice following i.p. injection with PBS, IL-1β (5 μg/kg), ketorolac (15 mg/kg)+IL-1β, or DCZ (100 μg/kg)+ketorolac+IL-1β. **(G)** Two-way repeated-measures ANOVA: main effects of time, treatment, and time×treatment interaction, p < 0.0001. n = 9 mice, paired design. **(A–I)** Error bars indicate mean ± SEM. Lines represent individual mice. Holm–Šídák post hoc comparisons: *p<0.05, **p<0.01, ***p<0.001, ****p<0.0001

